# Graph Laplacian Spectrum, topological properties, and high-order interactions of Structural Brain Networks are Subject-Specific, Repeatable but Highly Dependent on Graph Construction Scheme

**DOI:** 10.1101/2023.05.31.543029

**Authors:** Stavros I Dimitriadis

## Abstract

It has been proposed that the estimation of the normalized graph Laplacian over a brain network’s spectral decomposition can reveal the connectome harmonics (eigenvectors) corresponding to certain frequencies (eigenvalues). Here, I used test-retest dMRI data from the Human Connectome Project to explore the repeatability, and the influence of graph construction schemes on a) graph Laplacian spectrum, b) topological properties, c) high-order interactions (3,4-motifs,odd-cycles), and d) their associations on structural brain networks (SBN). Additionally, I investigated the performance of subject’s identification accuracy (brain fingerprinting) of the graph Laplacian spectrum, the topological properties, and the high-order interactions. Normalized Laplacian eigenvalues were found to be subject-specific and repeatable across the five graph construction schemes. The repeatability of connectome harmonics is lower than that of the Laplacian eigenvalues and shows a heavy dependency on the graph construction scheme. A repeatable relationship between specific topological properties of the SBN with the Laplacian spectrum was also revealed. The identification accuracy of normalized Laplacian eigenvalues was absolute (100%) across the graph construction schemes, while a similar performance was observed for a combination of topological properties of SBN (communities,3,4-motifs, odd-cycles) only for the 9m-OMST. Collectively, Laplacian spectrum, topological properties, and high-order interactions characterized uniquely SBN.

## 1. Introduction

The human brain can be modeled as a graph *G* = (*V, E*), comprising of nodes, V, representing brain regions and edges, E, referring to functional and anatomical strengths (Bullmore and Sporns, 2012). A high repertoire of network metrics has been adopted from social network analysis and applied to the analysis of human brain networks. Those metrics quantify different global and local properties of nodes such as the degree, the communication efficiency, etc (Boccaletti et al., 2006; Newman, 2003). Complementary to trivial network metrics, researchers have proposed a variety of qualitative measures for the examination of the global structure of brain networks (Atay et al., 2006; Banerjee and Jost, 2007; Banerjee, 2012; Varshney et al., 2011).

The eigenanalysis of the graph Laplacian operator over the structural brain network reveals a set of graph Laplacian eigenvectors and eigenvalues. The graph Laplacian eigenvectors, called connectome harmonics, is a set of frequency ordered harmonic patterns arising from the cortex and can be seen as a connectome extension of the well-known Fourier basis of a 1D signal to the 2D human brain network. These connectome harmonics reported a relationship between low-frequency harmonics (eigenvectors linked to smaller Laplacian eigenvalues) and the resting-state brain activity mainly from the default mode network (DMN) measured by functional magnetic resonance imaging (fMRI) recordings (Atasoy et al., 2016, 2018b).

The transformation of the original brain network to the normalized Laplacian matrix gives us the opportunity to estimate the Laplacian eigenvalues which refer to the global network structure (Banerjee, 2012; Chung, 1996). The advantage of the normalized Laplacian spectrum over unnormalized is that all the relevant eigenvalues range between 0 and up to a maximum of 2, which further enables the comparison of networks across modalities, cohorts, age groups, and even sizes (Banerjee, 2012).

Research studies have applied spectral graph theory to neural networks (Banerjee and Jost, 2007; Varshney et al., 2011), proposing network metrics tailored to the eigenvectors of the brain network, the node centrality (Bonacich, 2007, 1972; Page et al., 2001) and community detection methods (Fortunato, 2010; Harriger et al., 2012; Liang et al., 2011; Newman, 2006). The eigenvectors of a network are related to the local properties of a node over its neighborhood while the associated eigenvalues contain important information about the graph structure (Banerjee and Jost, 2007; Banerjee, 2012; McGraw and Menzinger, 2008; Vukadinović et al., 2002).

Many researchers have already applied the graph Laplacian to brain connectivity networks reporting the progression of neurodegenerative diseases (Raj et al., 2012), brain malformation (Wang et al., 2017), attention switching period in a cognitive task (Huang et al., 2018; Medaglia et al., 2018), macroscale coupling gradient between brain regions (Preti and Van De Ville, 2019), structure-function decoupling (Griffa et al., 2022), and an aberrant dynamic connectivity profile constrained by structural brain network in patients with concussion (Sihag et al., 2020). These studies focus on long-range and white-matter-based anatomical connectivity employing brain networks of sizes from a few tens up to a few hundred regions-of-interests (ROI) (Desikan et al., 2006; Destrieux et al., 2010). Recently, Atasoy et al. proposed an alternative framework for the application of graph Laplacian to the analysis of the human connectome. They combined assessment of local connectivity of the gray matter cortical structure captured from the magnetic resonance imaging (MRI) data with the assessment of long-range connectivity mediated via the white-matter thalamocortical fibers captured from the diffusion MRI (dMRI) data into a common anatomical network without the use of a template (Atasoy et al., 2017, 2016; Naze et al., 2021).

After many years of reductionism in science, researchers understood that no matter how accurate is our knowledge at the level of subsystem, we will miss the linear, and nonlinear interactions between the system components (Anderson, 1972). This is the reason, why we cannot fully explain the starting point of epileptic seizures just from the individual neurons of the human brain. Epilepsy is now conceptualized as a network disease (Lehnertz et al., 2023). Over the past decades, a variety of complex systems has been successfully described as networks whose interacting pairs of nodes are connected by links (Barabási,2011). Among these complex systems that are modelled as networks are the brain networks (Stam,2014). Breakthrough papers on networks were introduced to the scientific community close to millennium building upon earlier work in social network analysis, and mathematics (Watts and Strogatz,1998; Barabási and Albert, 1999). These pioneer studies triggered an exponential increment of published articles up to our days forming a new multidisciplinary field called Network Science.

Up to now, the majority of research on network science, and its applications on every discipline such as brain network analysis focused mostly on pairwise interactions (Battiston et al.,2020). However, interactions in general, and specifically in brain can often occur in groups of three or more nodes and cannot be described simply in terms of dyads (Sporns and Kotter,2004). Only recently, the community has devoted more attention to the high-order interactions introducing frameworks such as motifs, cliques, hypergraphs, simplicial complexes etc (Battiston et al., 2021).

Here, I will focus on exploring high-order coordinated structural pattern encoded with motifs. Motifs are small recurrent subgraphs with specific connectivity pattern that are considered as a high-order structural signature of the underlying network’s function (Milo et al., 2002). Motifs allow to extract additional information on the properties of an interaction, while can be described of each edges (1-interactions) between vertices that appear to be statistically significant in the network. However, a drawback of the motifs’ research is that the whole set of possible motifs to explore grows exponentially as the number of nodes involved in the whole analysis. Considering high-order interactions in human brain neuroimaging can help us understand many timeless mysteries of the human functionality such as consciousness (Herzog et al., 2024).

The analysis of the relation between the spectrum of a graph, i.e., the eigenvalues of its adjacency matrix, and the structural properties of a network is the main goal of spectral graph theory. Graph Signal Processing (GSP) is a special area in signal processing based on spectral graph theory where the data possess an intrinsic graph structure here a SBN. GSP extends graph theoretical approaches, providing an elegant and concrete mathematical framework to describe brain function as signal diffusion through the structural connectivity (Lioi et al., 2021; Abdelnour et al., 2014). At the centre of GSP lies the graph Laplacian matrix and its decomposition into graph harmonics called eigenvectors or “gradients” (Margulies et al., 2016), reflecting orthogonal spatial patterns of a signal in the network, while every harmonic is associated to an eigenvalue reflecting its graph frequency (Chung, 1996; Deslauriers-Gauthier et al., 2020; Luppi et al., 2020). The harmonic decomposition of the graph Laplacian matrix is similar to the Fourier transform of a signal. Laplacian harmonics form the set of Fourier basis that describes how brain signals would reflect in the structural brain networks, a transformation that link structural graph topology to functional synchrony (Müller et al., 2017). Each harmonic is associated with an eigenvalue, and conventionally harmonics are sorted by their ascending eigenvalues.

Based on the aforementioned definitions, it is natural to employ the harmonic “eigenspectrum” as the organizing principle that can link the integration and segregation which are the two ends of a continuum from the synchrony to asynchrony (Deco et al., 2015; Sipes et al., 2024). Integrative and segregative harmonics occupy the ends of the continuum while degenerate harmonics are in the middle of the continuum (Sipes et al., 2024).

In the present study, the general theme was then, firstly, to compute the eigenvalues of such matrices, and secondly, to relate the eigenvalues to structural properties of graphs such as: the synchronizability, the Laplacian energy, the relative frequency (RF), the number of communities (modules), the bipartiteness, the high-order interactions such as the motifs’ distribution and the distribution of odd-cycles. In the present study, I investigated the repeatability: a) of the Laplacian eigenvalue spectrum, b) of the Laplacian eigenvectors (connectome harmonics) of the structural brain networks derived from diffusion magnetic resonance imaging data (dMRI), c) of the structural properties of SBN including high-interactions investigated via motifs, and d) of the association between Laplacian eigenvalue spectrum and the structural properties of the dMRI-based brain networks. I analysed the test-retest MRI and diffusion-MRI data set from the multimodal neuroimaging database of the Human Connectome Project (HCP) (Glasser et al., 2013; S N Sotiropoulos et al., 2013; Van Essen et al., 2013).

The main aims of the present study were unique in human brain network neuroscience, and especially in dMRI-based SBN. It is the very first time in the literature according to my knowledge, that the Laplacian spectrum, the relevant properties of the SBD, and especially the high-order network interactions, and their association are studying together in structural brain networks. On the top, the whole investigation includes brain fingerprinting performance of Laplacian spectrum, and of the adopted structural properties of the SBN. Additionally, the current study took the advantage of this valuable open test-retest study to explore the repeatability of my findings (Dimitriadis et al., 2021; Messaritaki et al., 2019). As in our previous studies, I demonstrated how all these observations were influenced by the different graph construction schemes and the alternative network weighting schemes (Qi et al., 2015).

I constructed structural brain networks from this test–retest diffusion MRI scan data from the Human Connectome Project (HCP) using the b = 2000 s/mm^2^ data and selecting five out of seven most reproducible graph-construction schemes as derived from our previous study on the same data (Messaritaki et al., 2019a).

The major aim of this study was to investigate the associations of graph Laplacian spectrum with topological descriptors of the architecture of SBN. Simultaneously, I explored the repeatability of these observations and how could be influenced by alternative graph construction schemes. Finally, I examined how graph Laplacian spectrum and relevant brain network descriptors of SBN can produce a concrete system-level fingerprint of brain networks following a brain fingerprinting approach (de Lange et al., 2014,2016).

The rest of this manuscript is organized as follows: Section 2 (Methods) describes briefly the cohort, the processing of dMRI test-retest dataset, the graph construction schemes, the estimation of graph topological descriptors and their associations with graph Laplacian spectrum. Section 3 (Results) reports our findings in terms of repeatable normalized Laplacian eigenvalues and eigenvectors and their subject specificity under the brain fingerprinting framework. Section 4 (Discussion) summarises the major contribution of my study explaining its advantages, limitations, and possible future directions.

## 2. Methods

All analyses were performed using MATLAB (2019a; The Mathworks, Inc., MA).

### 2.1. Data

My study adopted the test-retest MRI and diffusion-MRI dataset from the large multimodal neuroimaging database of the Human Connectome Project (HCP) (Glasser et al., 2013; S N Sotiropoulos et al., 2013; Stamatios N Sotiropoulos et al., 2013; Van Essen et al., 2013). The cohort used in my study consists of 37 subjects which were scanned twice with a time interval between the scans ranging between 1.5 and 11 months. The age range of the participants was 22–41 years. It should be noted that the test-retest time interval is shorter than the expected time over which maturation-induced structural changes can be measured with the diffusion MRI (dMRI) experiment reported in this study.

The diffusion-weighted images (DWIs) had a resolution of (1.25×1.25×1.25) mm^3^ and were acquired at three different diffusion weightings (*b*-values: 1000 s/mm^2^, 2000 s/mm^2^ and 3000 s/mm^2^) across 90 gradient orientations for each *b*-value. The HCP acquisition details and pre-processing are described in (Feinberg et al., 2010; Glasser et al., 2013; Moeller et al., 2010; Setsompop et al., 2012; S N Sotiropoulos et al., 2013; Stamatios N Sotiropoulos et al., 2013; Xu et al., 2012).

### 2.2. Tractography

In our previous studies employing the same dataset, we performed tractography using the constrained spherical deconvolution (CSD) algorithm (Dimitriadis et al., 2021; Messaritaki et al., 2019). Here, I performed tractography with a probabilistic, anatomically constrained streamline tractography using MRtrix (Tournier et al., 2019), employing the iFOD2 (Second-order Integration over Fiber Orientation Distributions) algorithm (Smith et al., 2012, 2015; Tournier et al., 2010). The selected parameters of the algorithm were : a) the minimum and maximum streamline lengths were ranged between 30 mm and 250mm, b) the maximum angle between successive steps was defined to 50°, and c) the FOD amplitude cut-off was set-up to 0.06.

A total amount of two million streamlines were generated for each participant, and in both scans, with the seed points to be set on the interface between grey matter and white matter. I performed a visual inspection of the tractograms as a way to secure that the white matter was covered, and streamlines didn’t out of the white matter space.

IFOD2 was applied to DWI data acquired with b=2000 s/mm^2^.

### 2.3. Graph generation

Different experimental protocols and researcher’s methodological choices can alter the final structural brain network (SBN) (Qi et al., 2015). SBN and the extracted topological network measures can vary remarkably across different MRI gradient schemes and orientation models (Zalesky et al. 2010). Bastiani et al. (2012) reported that CSD generates a higher edge density and global efficiency, and lower small-worldness than diffusion tensor imaging (DTI). Tractography algorithms can also affect the derived SBN. Bastiani et al. (2012) showed that probabilistic methods lead to higher edge density than the deterministic ones. Global tractography also generates a higher edge density than local tractography, likely due to a higher number of longer connections. Moreover, many other factors such as the initial seed point region, the number of seed points, and the tracking termination criteria also affect the extracted SBN and the related network measures. Additionally, alternative network weighting schemes have been proposed for the construction of weighted SBN (Qi et al., 2015). For a nice review, an interested reader can check Qi et al., (2015). In the present study, I will focus only on how the different graph construction schemes can alter the topology of SBN.

#### 2.3.1. Parcellation, and Node definition

As in our previous studies, I adopted the Automated Anatomical Labeling (AAL) atlas (Tzourio-Mazoyer et al., 2002) to define 90 cortical and subcortical areas (45 areas per hemisphere) as nodes of the constructed structural brain graphs. Structural brain networks (SBN) were generated for each participant, scan and for each edge weight (see section 2.3.2) using ExploreDTI-4.8.6 (Leemans et al., 2009).

#### 2.3.2. Alternative Network Weighting Schemes

SBN are originally weighted and capture the information of connectivity attributes and strengths. Different weighting schemes can be employed for this scope. For example, the most straightforward scheme might be to utilize the number of fibers (or streamlines) connecting a pair of cortical regions as the weight, NS or the streamline density, (SLD), which is defined as the number of streamlines between two brain areas (nodes) divided by the mean volume of the two brain areas (Buchanan et al., 2014). Furthermore, the weight of the streamline density can be corrected by streamline length to generate WDL (Hagmann et al., 2008). A few important scalar metrics are fractional anisotropy (FA), Radial Diffusivity (RD) and Mean Diffusivity (MD) that have been interpreted as changes in the integrity of white matter microstructure for brain diseases and age-related morbidities, where the dMRI data are modelled by a locally anisotropic diffusion process (Jones et al., 2013).

Correspondingly, these scalar metrics can be useful for weighting connectivity. For instance, the weight of FA is adopted in a study by Buchanan et al. (2014). Microstructural white-matter properties (e.g., general fractional anisotropy (GFA), Intra-Cellular Volume Fraction (ICVF), Orientation Dispersion Index (ODI) (Lemkaddem et al., 2014) and physical distance properties (e.g., stream-line length, Euclidean distance between the nodes) (Bassett et al., 2011) can be estimated from dMRI as alternative weighting metrics for constructing SBN.

The motivation of using a linear combination of various metrics as edge weights is that the integration of the brain’s properties are affected by more than one attribute of the white matter tracts. Previous studies have also employed various combinations of metrics as network weighting schemes. Moreover, the edge-weight in a SBN can be the combination of aforementioned various measures. For example, the weight can be a product of the weight of the FN and the weight of the mean FA along a fiber bundle connectivity a pair of brain areas (Zhang et al., 2011). Nigro et al. (2016) used the product of NS and FA to weigh the edges in a study of Parkinson’s patients, and Taylor et al. (2015) used a combination of NS and TL in a study of epilepsy patients.

In the present study, I weighted the edges of the SBN by adopting the five most repeatable graph-construction schemes revealed previously with the same dataset (Messaritaki et al., 2019b), which were based on alternative combinations of the nine metrics listed in Table 1 (see Section 2.3.4). The edge weights of every SBN were normalized to have a maximum edge weight of 1, while the elements in the main diagonal were set to zero.

**Table 1.** Metrics used in connectivity matrices.

| Metric | Abbreviation |
| --- | --- |
| Fractional anisotropy | FA |
| Mean diffusivity | MD |
| Radial diffusivity | RD |
| Number of streamlines | NS |
| Percentage of streamlines | PS |
| Streamline density | SLD |
| Tract volume | TV |
| Tract length | TL |
| Euclidean distance between nodes | ED |

#### 2.3.3 Integrated Edge-Weights

Each metric shown in Table 1 conveys different information regarding the tissue properties. We previously proposed an integrated edge weighting scheme combining the metric-based SBN under a data-driven whole-brain algorithm (Dimitriadis et al., 2017a,b,c). An integrated SBN was formed by the combination of the nine metric-based SBNs for every participant and scan session.

An orthogonal-minimal-spanning-tree (OMST) algorithm was applied to every metric-based SBN, selecting edges of both small and large weights that preserved the efficiency of brain regions at a minimal wiring cost. The overall algorithm with the OMST on its center down-weights the metrics with a higher global topological similarity and up-weights the dissimilar metrics enhancing the complementarity of topological information across the nine adopted metrics. More details on the OMST algorithm and its implementation can be found in our previous work (Dimitriadis et al., 2017b,c, 2021, 2017a) and the related code is freely available at https://github.com/stdimitr/multi-group-analysis-OMST-GDD.

#### 2.3.4 Graph Construction Schemes

I will briefly explain the five graph construction schemes used here as in our previous studies.

The first category includes SBN constructed via the data-driven algorithm (Dimitriadis et al., 2017b, 2017a, 2017c).

A) NS-OMST: apply the OMST filtering algorithm (Dimitriadis et al., 2017b, 2017a, 2017c) to the NS-weighted matrix.

B) 9m-OMST: Integrate all nine diffusion metrics (as originally reported in Dimitriadis et al. 2017b, see Table 2).

**Table 2.** Summary of the graph-construction schemes. (number of streamlines (NS), fractional anisotropy (FA), and mean diffusivity (MD), orthogonal minimal spanning trees (OMST))

| Abbreviation | Initial Edge Weights | Topology | Final Edge Weights | Symbol |
| --- | --- | --- | --- | --- |
| NS – OMST | NS | OMST | Unchanged | A |
| 9-m OMST | lin. comb. of all 9 metrics in <u>Table 1</u> | OMST | Unchanged | B |
| NS-thr | NS | keep highest-NS edges | Unchanged | C |
| NS-t/FA-w | NS | keep highest-NS edges | re-weight with FA | D |
| NS-t/MD-w | NS | keep highest-NS edges | re-weight with MD | E |

The second category includes SBNs with edges weighted by the number of streamlines (NS), fractional anisotropy (FA), and mean diffusivity (MD) with various combinations of applying absolute thresholding on one individual metric-based SBN while keeping the same sparsity as the 9m-OMST that showed the highest reproducibility (Messaritaki et al., 2019). Since, absolute thresholding cannot guarantee the connectedness of the network, I first applied a minimal spanning tree (MST) on the original SBN constructed by NS. Then, I applied an absolute threshold on the rest of the NS weights (excluding the ones constitutes the MST) defined such as to return a SBN with the same density as the one returned by the 9m-OMST. After the MST and the absolute thresholding steps, the topology was either kept as it was (C) or re-weighted with one of the remaining two metrics (D,E) (see Table 2).

C) NS-thr: MST plus keep the highest-NS edges to align the density to 9m-OMST

D) NS-t/FA-w: Threshold to keep the highest-NS edges, then reweight those edges with their FA.

E) NS-t/MD-w: Keep the highest-NS edges, then reweight those edges with their MD.

In previous studies, we ranked twenty-one graph construction schemes with similarities ranging from 0.99 to 0.42 (Table 3; Messaritaki et al., 2019; Dimitriadis et al., 2021). Here, I focused on the first five graph construction schemes with the highest topological similarity (Table 2).

**Table 3.** Group-mean λ_2_, group mean Eigengap and the number of communities based on eigen-difference and K-Means clustering over the first eigenvectors across subjects for every graph construction scheme. I underlined with bold, the p-values that showed significant differences compared to the surrogate-based p-values. (Letters from A to E refer to the five graph construction schemes defined in Table 2.

|  | A | B | C | D | E |
|--|---|---|---|---|---|
| $\lambda_2(\text{mean} \pm \text{std})$ | 0.67±0.04 | 0.70±0.04 | 0.41±0.28 | 0.23±0.05 | 0.18±0.04 |
| $\lambda_2^{\text{surr}}(\text{mean} \pm \text{std})$ | 0.39±0.04 | 0.40±0.05 | 0.39±0.21 | 0.21±0.04 | 0.16±0.04 |
|  | (p = **0.00015**) | (p = **0.00005**) | (p = 0.071) | (p = 0.064) | (p = 0.068) |
| Eigengap(mean±std) | 2.61±0.24 | 3.42±0.56 | 2.76±0.65 | 2.31±0.57 | 2.21±0.23 |
| Eigengap<sup>su</sup><sub>rr</sub>(mean±std) | 1.48±0.52 | 2.32±0.47 | 2.23±0.58 | 2.09±0.63 | 1.65±0.32 |
|  | (p = **0.0007**) | (p = **0.00038**) | (p = 0.072) | (p = 0.079) | (p = 0.071) |
| K-Means(mean±std) | 7.67±1.35 | 8.72±0.62 | 4.41±1.34 | 2.81±1.34 | 2.65±0.78 |
| K-Means<sup>surr</sup>(mean±std) | 3.45±1.45 | 4.23±1.42 | 3.82±1.23 | 2.10±1.56 | 1.56±0.48 |
|  | (p = **0.00019**) | (p = **0.00041**) | (p = 0.634) | (p = 0.712) | (p = 0.685) |

SBN built with 9m-OMST, NS-thr, NS-t/FA-w and NS-t/MD-w graph construction schemes share the same density but a different topology while SBN constructed with NS-thr, NS-t/FA-w and NS-t/MD-w graph construction schemes share the same topology with different edge weights.

Fig.1A illustrates the five SBNs from the first scan of the first subject.

**Fig. 1.**
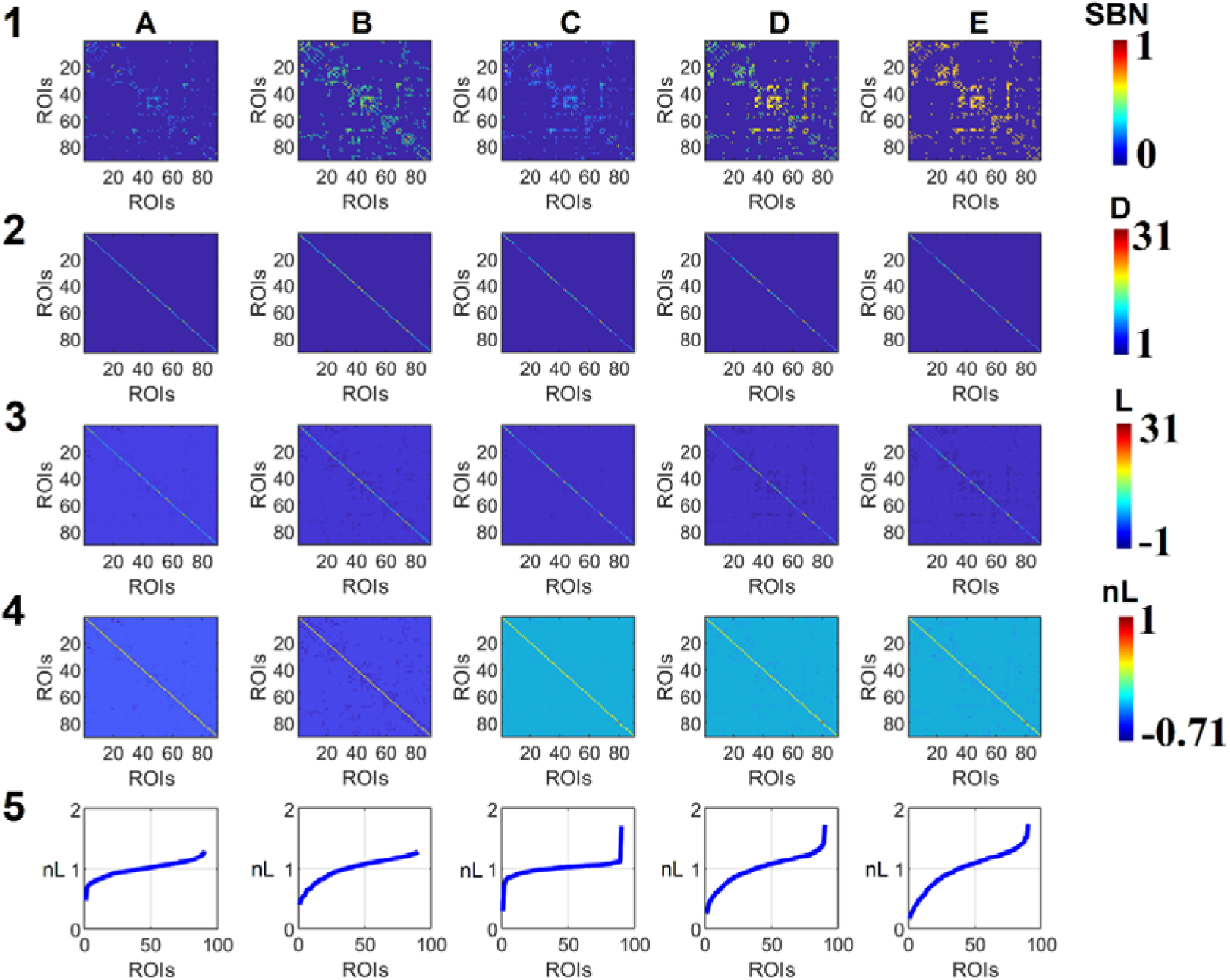
Illustration of preprocessing steps for the estimation of normalized Laplacian eigenvalues. The data are derived from the first scan of the first subject from the dMRI cohort. A-E in columns refer to the five graph construction schemes tabulated in Table 2. Numbers refer to the processing steps: 1. Original metric-based SBN for the five graph construction schemes as they are reported in Table 2. 2. The D degree matrices of the five SBN shown in A 3. The unnormalized Laplacian matrix L of the five SBN 4. The normalized Laplacian matrices nL of the five SBN 5. The normalized Laplacian eigenvalues (nL) linked to the five SBN

### 2.4 Laplacian Spectrum

#### 2.4.1 The normalized Laplacian matrix

In this paper, I considered the transformation of individual integrated SBNs from the five graph construction schemes to the normalized Laplacian matrix L. The Laplacian matrix L has the advantage that its eigenvalues range between 0 and 2, enabling the direct comparison of SBN across modalities, subjects, cohorts, and of different sizes (Chung, 1996).

The normalized Laplacian matrix is defined as:

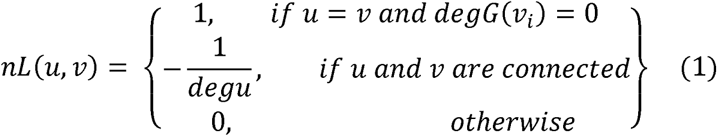

with u and v representing two nodes of the network or brain regions, L(u,v) the edge from node u to v and degu the degree of node u which is the total number of its connections. Fig.1 illustrates the processing steps needed from the original SBN up to the estimation of the normalized Laplacian spectrum given by the normalized Laplacian eigenvalues. Rows correspond to the processing steps of extracting the normalized Laplacian eigenvalues from SBN while columns refer to the five graph construction schemes.

The normalized Laplacian matrix can be also estimated and expressed from its relation with the adjacency matrix A as nL = I – *D*^-1/2^ × L × *D*^-1/2^ (Fig.1D) where the D is the degree matrix, where its diagonal elements encapsulate the degree of every node (Fig.1B), L = D - A is the unnormalized Laplacian matrix (Fig.1C), and A is the adjacency matrix. The eigenanalysis of the nL extracts a collection of eigenvalues λ for which a non-zero vector eigenvector v exists that satisfies the equation *Lv* = λ*v*.

Eigenvalues share important properties. The multiplicity of the eigenvalues equal to 0 (λ = 0) is equal to the number of connected modules (Chung, 1996). The largest eigenvalue is equal to or smaller than 2, sorting the range of eigenvalues as 0 ≤ λ1 ≤… ≤λ*n* ≤ 2 ( (Chung, 1996); Fig.1E).

#### 2.4.2 Repeatability of Laplacian Eigenvalues

I quantified the repeatability of Laplacian eigenvalues using Pearson’s correlation coefficient (Pcc) accompanied by the relevant p-value. The Pcc was estimated between Laplacian eigenvalues derived from the two scan-sessions from the same graph construction scheme (within graph construction scheme) and also between graph construction schemes (5×4/2 = 20 pairs) from the same or different scan session (between graph construction schemes).

I then estimated the group-mean Pcc across the cohort related to within-session and graph construction scheme and the group-mean Pcc linked to between-session and graph construction scheme. The Pcc values for the between-session approach were first averaged across the two scans for the 20 pairs and then across the 20 pairs of comparisons. Adopting a Wilcoxon Signed Rank-Sum test, I estimated the significance level between the two sets of subject-specific Pcc values on the subject level that will support at which degree the Laplacian eigenvalues are highly dependent on the graph construction scheme.

#### 2.4.3 Repeatability of Important nL-based properties

Network synchronizability of a variety of complex networks can be characterized by the ratio of the second smallest eigenvalue λ_2_ to the largest eigenvalue of the Laplacian matrix λ_n_ (Barahona and Pecora, 2002). So Synchronizability = λ_2_ / λ_n_.

The following formula (2)

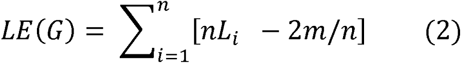

is called the Laplacian energy of the network G, where nL are the normalized Laplacian eigenvalues, m is the number of edges and n is the number of vertices (Hakimi-Nezhaad and Ashrafi, 2014).

The repeatability of the Synchronizability and Laplacian energy per graph construction scheme and between the two scans was quantified with the absolute difference between the two scans. The original values were compared with the surrogate ones per graph construction scheme and scans. A p-value is assigned to the original values by a direct comparison with ten thousand surrogate values.

I applied a Wilcoxon Signed Rank-Sum test for both Synchronizability and Laplacian Energy properties between the two scans.

#### 2.4.4 Laplacian Spectrum Properties

Many important dynamical network models can be formulated as a linear dynamical system which can be expressed by the following diffusion equation

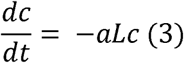

which is a continuous time version.

As I mentioned before, the Laplacian matrix of a network is expressed as L = D – A. The Laplacian matrix is symmetric in which diagonal components are all non-negative (representing node degrees) while the other components are all non-positive.

A ***Laplacian matrix*** of an undirected network has the following interesting properties:

1. At least one of its eigenvalues is zero.
2. All the other eigenvalues are either zero or positive.
3. The number of its zero eigenvalues corresponds to the number of connected components in the network.
4. If the network is connected, the dominant eigenvector is a homogeneity vector h=(11…1)^T^.
5. The smallest non-zero eigenvalue is called the *spectral gap* of the network, which determines how quickly the diffusion takes place on the network.

##### Smaller Eigenvalues

Laplacian eigenvalues and the relevant eigenvectors play an important role on the studying of multiple aspects of complex network structures like resistance distance, spanning trees and community structures (Newman,2006). According to Newman’s study, only the eigenvectors related to positive eigenvalues could contribute to the partitioning of the network and to the modularity. This practically means that the optimal graph partitioning could be achieved by selecting the number of communities/groups in a network to be equal with the number of positive eigenvalues plus 1. In the normalized Laplacian graph, the important role of guiding the spectral clustering of the network is supported by the smallest normalized Laplacian eigenvalues. The k eigenvectors that correspond to the K smallest eigenvalues of the normalized Laplacian graph create a n x K matrix (where the n refers to vertices of the graph and K to the eigenvectors). Clustering the row eigenvectors 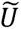 using K-Means will give us the number of communities of the network.

Given a graph Laplacian matrix L (here normalized L), spectral clustering proceeds to compute the eigenvalue decomposition of *L* = *UΣU^T^*. Then we choose the K smallest eigenvalues and extract the matrix 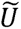 which contains the K columns of U corresponding to theses values. 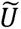 is of dimension n x K. Finally, I applied K-means algorithm to cluster the n row vectors of 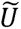. Node i is assigned to the cluster of the i^th^ row vector of 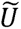

This is the famous **spectral clustering** with the following algorithmic steps:

A. Perform an eigenvalue decomposition of a graph Laplacian matrix, here the normalized Laplacian matrix nL :n*L* = *UΣU^T^*
B. Extract 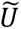 by taking the K columns of U corresponding to the K smallest eigenvalues
C. Cluster the row vectors of 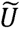 using K-means algorithm

Performing a K-Means clustering on the n vertices in the K-dimensional Euclidean space, one can reveal the communities of the graph. Based on the aforementioned properties, the smallest eigenvalues of the Laplacian spectrum reflect the modular organization of a network (Donetti, 2005; Fortunato, 2010; Shen and Cheng, 2010; Shi and Malik, 2000).

The spectral gap is the smallest non-zero eigenvalue of L, which corresponds to the largest non-zero eigenvalue of −αL and thus to the mode of the network state that shows the slowest exponential decay over time. The spectral gap’s value determines how quickly the diffusion takes place on the network. If the spectral gap is close to zero, this decay takes a very long time, resulting in slow diffusion. If the spectral gap is far above zero, the decay occurs quickly, and so does the diffusion. The larger the value of the first nonzero eigenvalue of *L* the faster the convergence of the diffusive process. In that sense, the spectral gap of the Laplacian matrix captures some topological aspects of the network, i.e., how well the nodes are connected to each other from a dynamical viewpoint. The spectral gap of a connected graph (or, the second smallest eigenvalue of a Laplacian matrix in general) is called the *algebraic connectivity* of a network.

Every eigenvector v_i_ informs us of a unique bisection of the nodes of a network assigning to each one a positive or negative value and the associated eigenvalues λ_i_ express the inverse diffusion time of this dichotomy to the stationary state. Smaller eigenvalues are indicative of longer diffusion times, revealing a larger proportion of inter-module connections and a smaller number of inter-module connections.

The λ_2_ eigenvalue provides the possible best division of nodes into two modules and it is called Fiedler value while the corresponding eigenvector is called Fiedler vector (Chung, 1996). One option that is proposed in the literature is the combination of divisions derived from all eigenvectors up to v_i_ to assign every node to i communities. A possible optimal number of communities can be defined by the largest eigen-difference (eigen-gap) between consecutive Laplacian eigenvalues (λ*_i_* _+ 1_ − λ_i_) (Cheng and Shen, 2010; Shi and Malik, 2000). In summary, small eigenvalues, their number, and the eigen-differences reflect important attributes of the modular structure of a network (Fig.2A).

**Fig. 2.**
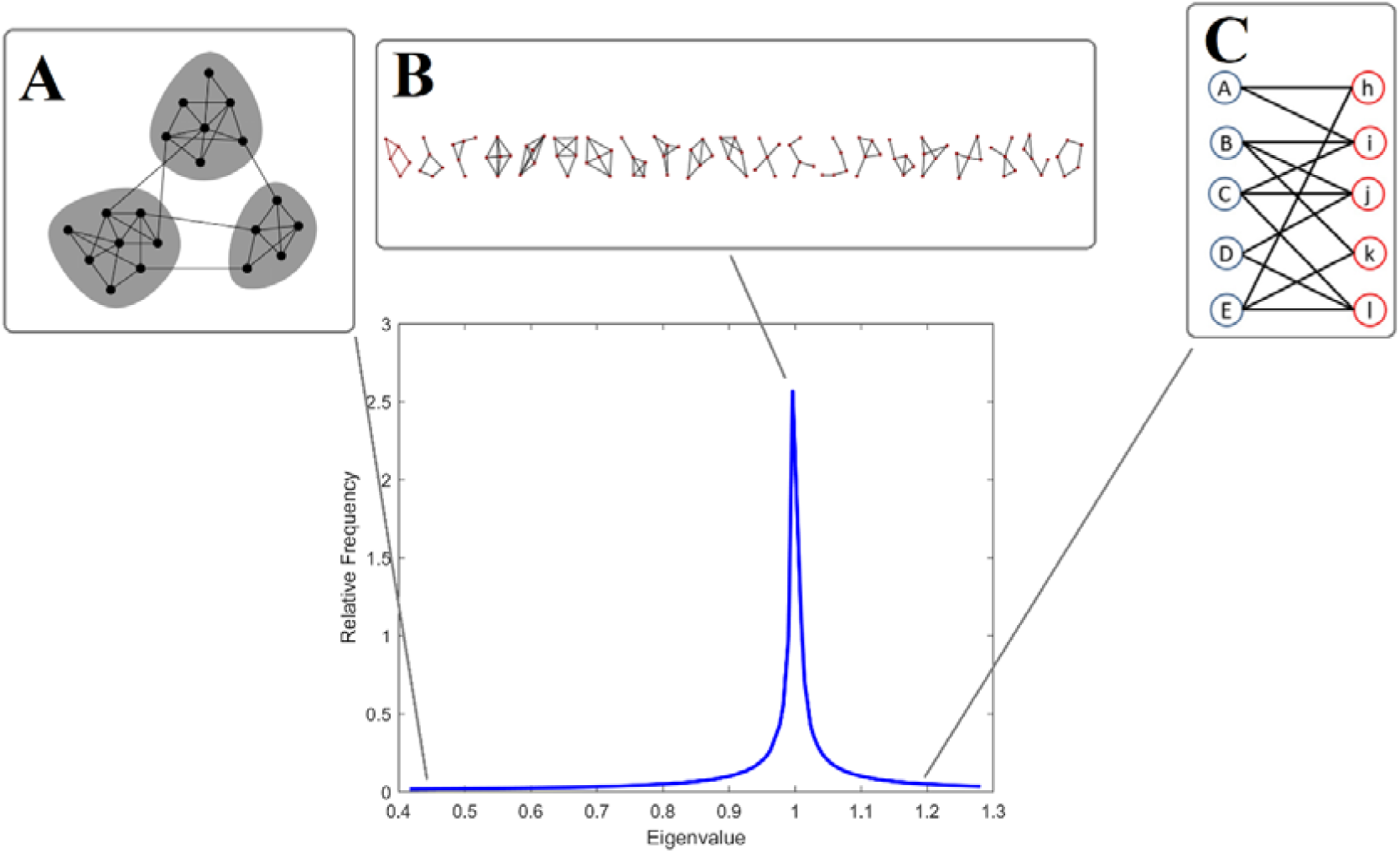
Topological properties of a network reflected in the Laplacian Spectrum. Relative Frequency (RF) is linked to peak at λ=1. (A) The first smaller Laplacian eigenvalues are indicative of stronger community structures (B) Recursive motifs in the complex network result in Laplacian eigenvalues of high multiplicities, revealing characteristic peaks in the Laplacian spectrum (λ=1) (C) The largest eigenvalue reflects the level of ‘bipartiteness’ of the most bipartite subgraph of the network which is alternatively linked to the total number of odd cyclic motifs of the network.

In the present study, I compared the methods of eigen-gap Laplacian differences with the K-means clustering applied over the K eigenvectors that correspond to the K smallest eigenvalues of the normalized Laplacian graph. The second approach creates a n x K matrix (where the n refers to vertices of the graph and K to the eigenvectors). The total sum of the normalized Laplacian eigenvalues in my study equals to 90. I defined the smallest eigenvalues, the first ones where their sum divided by the total sum overcomes the 10%. I ran the K-Means clustering 50 times on the n vertices in the K-dimensional Euclidean space integrating the findings to avoid the influence of the random initializations of the K-Means algorithm. I adopted mutual information (MI) as in our previous study (Dimitriadis et al., 2021) to measure the similarity of graph partitions per subject and graph construction scheme between the two scans. The outcome of graph partitions affiliations with both the methods is compared with the outcome of the best partition observed in our exploratory analysis with a high number of graph partition algorithms applied in the same dataset (Dimitriadis et al., 2021).

I estimated group-mean λ_2_ (Fiedler value) and group-mean number of communities defined by the eigen-difference of Laplacian eigenvalues (Eigengap method) and by the K-Means clustering applied over the first eigenvectors. These group-means were averaged first across scans and then across subjects. Group-mean λ_2_ and the number of communities defined by the two methods were compared with the surrogate number of communities. To quantify the similarity of graph communities between the two scans (repeatability) per graph construction scheme and in both methods, I employed MI as a proper measure. The original MI values were also compared with surrogate MI values for both methods adopting a Wilcoxon Rank-Sum test.

##### Medium Eigenvalues

Network motifs are statistically significant recurrent subgraphs. All networks like brain networks, biological, social and technological networks can be represented as graphs, which include a large variety of subgraphs. Practically, network motifs are repeatable sub-graphs that are defined by a specific pattern of interactions between vertices. They may also reflect a framework supporting particular functions to achieved in an efficient way (Sporns, and Kötter, 2004). For that reason, the motifs are of high importance to reveal the structural principles of complex networks reflecting their functional properties (Fig.2B). Motifs are characterized by their size that equals the number of vertices and by the repertoire of possible alternative ways that nodes are connected. By defining the number of the studying vertices M, I enumerated exhaustively the frequency of every single motif of size M across its structural connectivity variance. The outcome of this procedure gives the motif frequency spectra for structural motifs of size M. Usually, the size M is restricted within the range of [3 - 5] due to the computational power needed to enumerate exhaustively the motif frequency spectra of a network with a large number of vertices e.g. a few hundreds. Fig.3.A illustrates the repertoire of 2,3,4 motifs for an undirected graph.

**Fig. 3.**
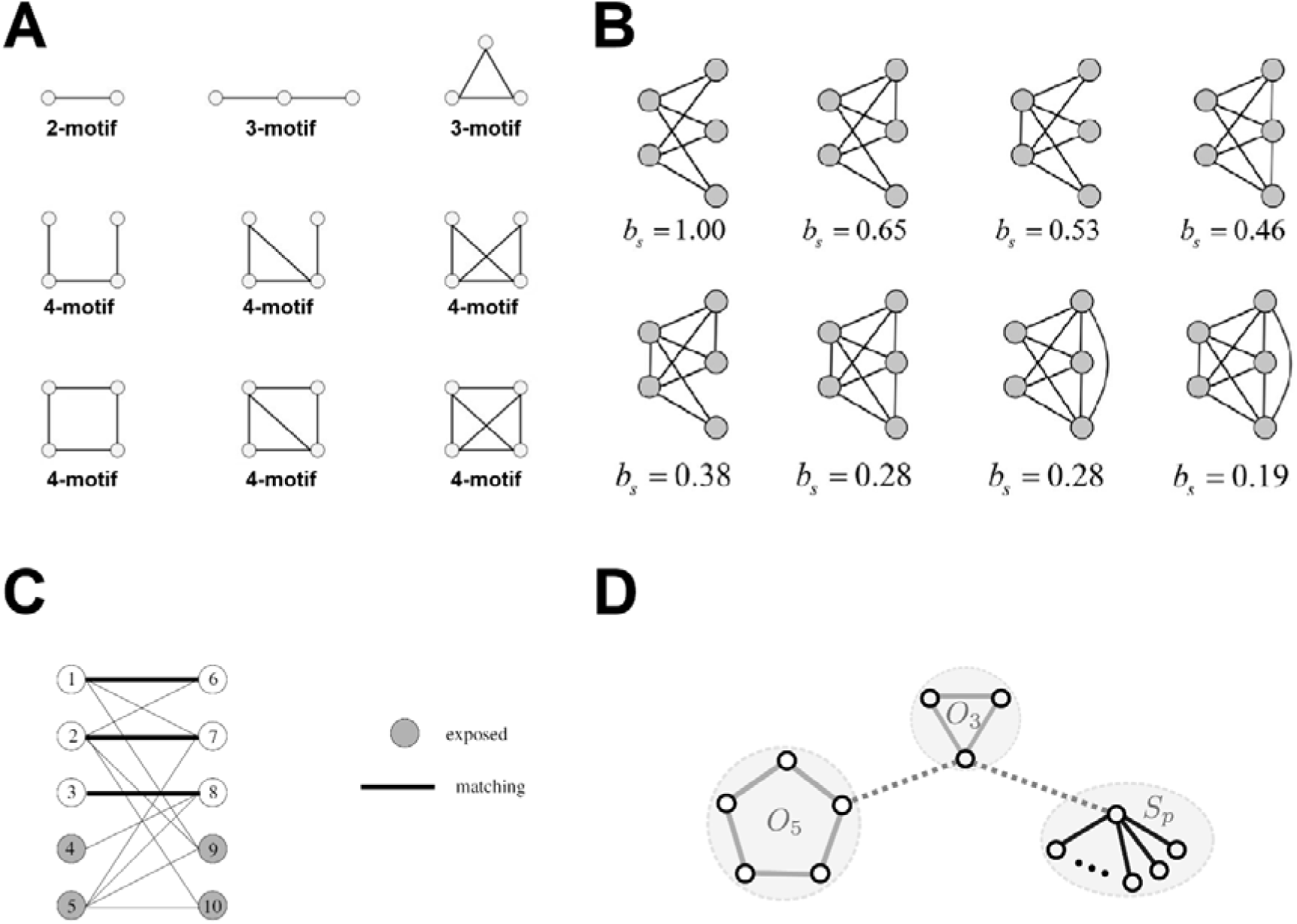
Motifs, bipartivitiy, odd cycles, and stars (Inspired and redesigned by Fig.8 in Estrada, 2021, Fig.3A in Biswas et al., 2021 and Fig.1 in lecture notes from Goemans, 2017). (A) The repertoire of 2-3-4 structural motifs in an undirected graph (B) Illustration of the monotonically change in the bipartivity index b_s_ with the increase in the number of ‘frustrated’ edges in a complete bipartite graph (C) The edges (1 - 6), (2 - 7) and (3 - 8) form a matching. Vertices 4, 5, 9 and 10 are exposed. (D) An example of an optimal decomposition of a motif *H* into odd cycles and stars. O3 refers to an odd-cycle of length = 3 that connects three nodes while O5 denotes an odd-cycle of length = 5 that connects five nodes.

It is well studied that repeated duplications and additions of nodes and motifs in the construction of a network leave traces in the network’s Laplacian spectrum (Banerjee and Jost, 2009, 2008). In a network, for example, two nodes with a similar connectivity pattern will increase the eigenvalue λ=1 of the spectrum (Banerjee and Jost, 2008). Duplication of edge motifs, for example, duplication of two connected nodes n_1_ and n_2_ has been shown to produce symmetrical eigenvalues around 1 with Laplacian eigenvalues 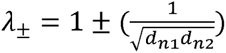, with *d*_n_ being the degree of node *n*. To better understand the relationship between motifs and eigenvalues, I give the following examples. An inclusion of a new triangle motif to a network results in the addition of an eigenvalue λ = 1.5 to the spectrum (Banerjee and Jost, 2008a). The joining or duplication of a motif in a network produces specific eigenvalues in the spectrum and repetition of these processes result in characteristic aggregated eigenvalues observed as peaks of the Laplacian spectrum. For that reason, the eigenvalues with high multiplicities e.g. high peak at λ = 1 or eigenvalues at equal distances around 1 are indicative of a local organization as a consequence of the presence of recursive motifs in the network (Fig.2B).

In the present study, I exhaustively quantified the 3,4-motifs across subjects, scans, and in the five graph construction schemes. The repertoire and the topology of structural 3,4-motifs is demonstrated in Fig.3A (Sporns, O., and Kötter,2004). A motif is a subnetwork consisting of N nodes and at least (N − 1) edges linking the nodes in a path. Network motifs are simple building blocks that characterized the complexity of information transfer within a network across many fields of science and their distributions deviates from those of random networks (Miro et al., 2002). I quantified the total number of structural 3,4-motifs in the network and also the motif frequency of occurrence around an individual node which is known as the motif fingerprint of that node (Figs7,8). The estimation of the motifs has been realized with proper functions of the brain connectivity toolbox (Rubinov and Sporns,2010).

Patterns of local connectivity are quantified by network motifs while simple measures of segregation are defined based on the total number of triangles in the whole network (global level). A high number of triangles implies strong segregation. In the local level, the fraction of triangles around a node is known as the *clustering coefficient* and is equivalent to the fraction of the node’s neighbors that are also neighbors of each other (Watts and Strogatz, 1998). I estimated the nodal weighted clustering coefficient and I correlated (using Pearson’s correlation coefficient) it to each of the two global 3-motif frequencies represented in Fig.7 (Rubinon and Sporns,2010). I followed the same approach per graph construction scheme. This approach will reveal the relationship between a segregation network metric (weighted clustering coefficient) and patterns of local connectivity (motifs).

A recent study proposed a measure to capture the global symmetry of a network and showed in both empirical and network models that the main peak in the Laplacian spectrum (λ = 1) is related to network elements that exhibit similar wiring patterns. The global symmetry of a network is quantified by the Matching Index (MIN) (Hilgetag et al.,2002) that expressed the similarity between nodes *i* and *j* and is inferred from the overlap between their connectivity pattern. De Lange et al., (2016) showed that global symmetry shaped neural spectra and the overlap in the wiring pattern of brain regions measured with MIN can explain the large central peak observed in spectra of neural networks (λ = 1). To reveal a link between the peak(s) of the Laplacian spectrum and the total number of 3,4-motifs, I adopted a multi-linear regression analysis per graph construction scheme between the relative frequency (RF) linked to peak at λ = 1 and the total amount of every possible structural 3 or 4 motif plus the Matching Index (MIN) with (Fig.2).

In summary, I estimated the group-mean relative frequency (RF) linked to peak around one (λ = 1) across subjects for every graph construction scheme, at first averaged between scans. The original RF values were compared with the surrogate ones adopting a Wilcoxon Rank-Sum test. As I aforementioned, I applied a multi-linear regression analysis between the RF and the total amount of each of the 3,4-motif and the MIN across the SBN independently per scan. I estimated Pearson’s correlation coefficient between the nodal weighted clustering coefficient and the nodal 3-motifs distribution across graph construction schemes. Complementary, I estimated Pearson’s correlation coefficient (Pcc) in a pairwise fashion between nodal motif frequency of occurrence across the five graph construction schemes. I followed this approach independently per subject, scan and for each of the two 3-motifs and six 4-motifs. These Pcc correlations were averaged across scans first and afterwards across subjects. To compare my findings with those present in de Lange et al., (2016), I estimated the Pearson’s correlation coefficient between the MIN and the RF per graph construction scheme and scan and I presented the mean across graph construction schemes averaged across scans.

#### Largest Eigenvalues

The largest eigenvalue of the Laplacian spectrum informs us of the level of ‘bipartiteness’ of the most bipartite subpart of the network, which is closely related to the number of odd cyclic motifs in the network (Bauer and Jost, 2009). A subnetwork is fully bipartite when its nodes can be divided into two groups where nodes of the same group are not connected.

A graph G = (V;E) is bipartite if the vertex set V can be partitioned into two sets A and B (the bipartition) such that no edge in E has both endpoints in the same set of the bipartition. A matching *M* ⊆ *E* is a collection of edges such that every vertex of V is incident to at most one edge of M. If a vertex v has no edge of M incident to it then v is said to be exposed (or unmatched). A matching is perfect if no vertex is exposed; in other words, a matching is perfect if its cardinality is equal to |*A*| = |*B*|. Fig.3C illustrates an example of perfect matchings and exposed edges.

The ‘bipartiteness’ is directly linked to the total number of odd cycle motifs in a network (Fig.2C & Fig.3D). Here, I also estimated the bipartitenes of the SBN with the following bipartivity index b_s_ (Estrada, 2022)

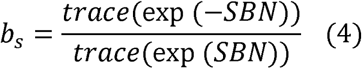

The bipartivity index b_s_ equals to 1 for a complete bipartite network while it changes monotonically with the increase of the number of edges “frustrating” the bipartition. The edges that if removed the network becomes bipartite are called frustrated. Such frustrated edges are shown in Fig.3B. One can see how the bipartivity index b_s_ changes monotonically with the increase in the number of ‘frustrated’ edges in a complete bipartite graph (Fig. 3B).

A motif H can be decomposed into a set of disjoint small cycles and stars and this decomposition is valid if all vertices of a motif H belong to either a star or an odd cycle in the set (Fig.3D). A star is a subnetwork type where only a central node is connected with the rest of the nodes while an odd cycle is a subnetwork with an odd number of vertices that are connected between each other in a circular way (Fig.3D). The degree of every node in an odd cycle is 2. It is important to underline here that bipartite graphs (*b_s_* = 1) do not contain odd length cycles, or graphs with odd length cycles are not bipartite (*b_s_* ≠ 1). If a graph is bipartite it doesn’t contain any odd length cycles, but, if a graph is non-bipartite it surely contains at least one odd length cycle.

I estimated the group-mean largest eigenvalue λ_n_ per graph construction scheme and compared it with the surrogate ones. The group-mean largest eigenvalue λ_n_ was at first averaged per scan across the graph construction schemes. Here, I estimated odd-cycles of length = 3,5 and 7 in an exhaustive way. In a similar way, I estimated the group-mean number of odd-cycles of length = 3,5,7 and the group-mean b_s_ per graph construction scheme and compared it with the surrogate ones. Both measurements were first averaged between scans.

In summary, I estimated the group-averaged of bipartivity index b_s_, of largest eigenvalue λn and of exhaustive estimation of odd-cycles of various lengths across subjects for every graph construction scheme, at first averaged between scans. The original values of bs, λn and odd-cycles were compared with the surrogate ones by adopting a Wilcoxon Rank-Sum test. Complementary, I applied a multi-linear regression analysis between the largest eigenvalue λn and the bs plus the total number of odd-cycles of length 3, 5 and 7 across the SBN independently per scan.

#### 2.4.5 Repeatability of Laplacian Eigenvectors

Ι estimated the repeatability of Laplacian eigenvectors (connectome harmonics) per graph construction scheme between the two scans by adopting the signum function (D^LS^).

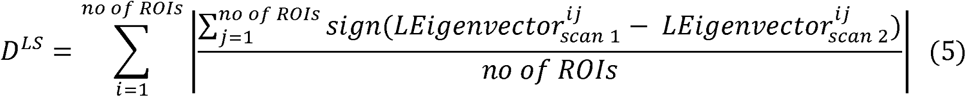

where no of ROIs denotes the number of brain areas while the first sum runs across Laplacian eigenvectors and the second sum across its vector size that equals the no of ROIs. The original group-mean D^LS^ values for every graph construction scheme was compared with the surrogate ones.

#### 2.4.6 Integrative, segregative, and degenerate harmonics of SBN

A recent study on SBN introduced a new framework that placed integration and segregation on the end of a continuum of structural connectivity graph Laplacian harmonics via the presentation of a gap-spectrum (Sipes et al., 2024). Gap-spectrum is estimated over sorted eigenvalues and naturally partitions the harmonics into three distinct regimes, the integrative harmonics that have low eigenvalues with high spectral gaps, the segregative harmonics that have high eigenvalues and high spectral gaps, and the “degenerate” harmonics that have intermediate eigenvalues but with low spectral gaps.

Gap-spectrum was defined as the derivative of the ascending eigenvalues accompanied with their index and presents a measure of harmonic degeneracy. The authors first fit an order 10 spline with 3 knots to the structural connectivity Laplacian eigenvalues to smooth the gap-spectrum due to the amplification of noise caused by derivative. Then, they computed the first analytical spline derivative as the gap-spectrum. They hypothesized that the first-order gap-spectrum (measuring harmonic degeneracy) would be related to various properties of harmonics. The implementation of the algorithm is presented on an open repository (https://github.com/Raj-Lab-UCSF/IntDegSeg/tree/main). The algorithm estimates the two main gaps that separate the Laplacian spectrum into the three regimes.

In my study, I estimated the gap-spectrum and the relevant integrative, segregative and degenerate harmonics on individual SBN and not on the consensus SBN (average SBN). The reason of this choice is to investigate the repeatability of these three regimes across subjects and graph construction scheme. Additionally, the estimation of a consensus SBN (averaging SBN across subjects) is impossible due to the individual network topology which involves weighted edges on subject-specific pairs of ROIs not consistent across the cohort. Averaging can be solely realized on fully-weighted brain networks that don’t really exist in any neuroimaging modality which is an old-fashioned methodology that made many assumptions while destroying any individual network topology.

The repeatability of individual three regional harmonics across scans was quantified with Embedded Laplacian Discrepancy (ELD), a newly introduced metric for Multiscale Graph Comparison of different size (Tam and Dunson, 2023). For that purpose, I introduced for the very first time on brain networks the use of a new metric tailored to Laplacian eigenvectors and eigenvalues for comparing graphs of different size, a property that is important in my study. There is no restriction on the consistency of graph-spectrum across scans and this is the main purpose of adopting such a metric that can compare the three defined harmonics of different size across scans.

The authors proposed the **Embedded Laplacian Discrepancy (ELD)** as a simple and fast approach to compare graphs (of potentially different sizes) based on the similarity of the graphs’ community structures. The ELD represents graphs as point clouds in a common, low-dimensional space, on which a natural Wasserstein-based distance can be efficiently computed in a multiscale way. A main challenge in comparing graphs through any eigenvector-based approaches is the potential ambiguity that could arise due to sign-flips and basis symmetries. To overcome this potential limitation, the ELD leverages a simple symmetrization trick to bypass any sign ambiguities. The ELD becomes a nice metric that encapsulates many interesting properties like invariance to graph isomorphism and invariance to signs configurations.

The theory behind ELD definition is based on the seminal paper (Belkin and Niyogi, 2003) where they showed that Laplacian decomposition provides a natural Laplacian - spectral embedding of a graph’s vertices in Euclidean space. This approach starts with the selection of the first *K Eigenvectors that corresponds to the first K eigenvalues of nL* (where *k* ∈ N^+^ is a hyperparameter), and continues with the employment of these eigenvectors as Euclidean coordinates for embedding the vertices in a common Euclidean space for comparisons. In other words, the *i*^th^ entry of the K^th^ eigenvector v_K_(*i*) provides the K^th^ coordinate of vertex *i*.

The Algorithmic steps of ELD estimation are the following:

1. Compute the top K Laplacian eigenvectors of the two graphs under comparison.
2. Use the entries of the K Laplacian eigenvectors as well as their flipped counterpart (for symmetrization) to represent the nodes of the two graphs as two point clouds in a common K-dimensional Euclidean space.
3. Compute the 1-dimensional Wasserstein distance of the point clouds along each of the K canonical Euclidean axes and average them.

Given the *r*^th^ eigenvector 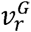 of a graph *G* with *n* nodes, the authors present it a one-dimensional empirical measure

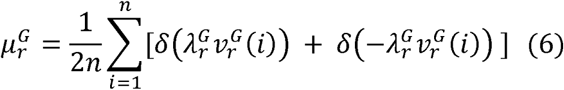

where *δ* is the Dirac measure and 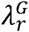 is the *r*^th^ eigenvalue. The negative (-) of the eigenvector is also included to symmetrize the embedding. In Section 4 of (Tam and Dunson, 2023), one can read how the embeddings are invariant to sign-flips of eigenvectors. I employed *μ* (eq. (4)) and analogously *ν* to define the empirical measures associated with the eigenvectors of the two different graphs.

The ELD is finally defined as. Consider two graphs *G*_1_ = (*V*_1_, *E*_1_,*w*) ∈ G and *G*_2_ = (*V*_2_, *E*_2_,*w*_2_) ∈ G, with sizes *n*_1_ = |*V*_1_ | and *n*_2_ = |*V*_2_ | and Laplacians 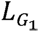 and 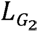,respectively. Without loss of generality we assume *n*1 ≤ *n*2. Given a dimension hyperparameter K ≤ *n*1, define the embedded Laplacian discrepancy as

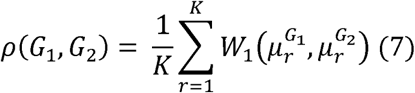

Here, K refers to the min(n_1_,n_2_) where n_1_,n_2_ are the number of eigenvalues within every region of the spectrum defined by the gap-spectrum approach (Sipes et al., 2024).

ELD is estimated between brain scans independently for the three regimes and graph construction scheme. I compared original ELD values with the surrogate ones independently for the three regimes and across graph construction schemes. I applied a Wilcoxon Rank-Sum test between the three regimes of harmonics in a pairwise fashion (3 pairs) and within every graph construction scheme and also per regime of harmonics across the graph construction schemes (5×4/2 = 10 pairs).

#### 2.4.7 Laplacian Spectrum’s Convolution

We further processes the Laplacian spectrum not as the collection of the eigenvalues λ_j_, but their convolution with a smoothing kernel, here a Gaussian, described by the following formula

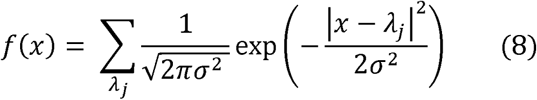

The particular smoothing value σ was set to 0.015. A discrete smoothed spectrum was used in which f had steps of 0.001. Furthermore, the distribution was normalized such that the total eigenvalue frequency was one. Relative Frequency (RF) linked to peak at λ=1 is estimated over the Laplacian spectrum after convolved with the above formula (Fig.2).

### 2.5 Brain Fingerprinting

As I have already mentioned, the human brain can be modelled as a network composed of brain regions (nodes) defined anatomically by a predefined brain atlas which are interconnected by two types of links or edges (de Reus and van den Heuvel, 2013). The structural connections can represent any attribute of white matter tracts assessed by dMRI leading to the structural connectome (Sporns et al., 2005). The functional connections represent statistical interdependencies between pairs of brain regions’ signals while subjects are either at rest or performing a task leading to the functional connectome (Friston, 2011). Structural and functional brain connectomics have been proven useful in mapping structural and functional properties between brain regions in large populations, but simultaneously in exploring the association between individual connectome features (connectomics) and clinical, behavioural and genetic profiles (Fornito et al., 2019; Sareen et al., 2020).

Fingerprinting is taking an ink impression of someone’s **fingerprints** for the purpose of identification. Brain fingerprinting was first used as an objective, scientific method to detect concealed information stored in the brain by measuring electroencephalographic (EEG) brain activity elicited non-invasively by sensors placed over the scalp (Farwell, 2012). Scientists present salient details about a crime or investigated situation as a way to place the subject to a crime scene via the release of a specific brainwave patter called the P300-MERMER. In neuroimaging, **the determination of individual uniqueness of brain activity or connectivity** is known as ‘brain fingerprinting’. Especially in cases where researchers employed brain connectome, it is called ‘brain connectome fingerprinting’ (Amico and Goñi, 2018; Finn et al., 2015; Miranda-Dominguez et al., 2014). Brain connectome fingerprinting is a new influential research field in brain connectomics that paves the way of extracting individual features from structural and functional connectomes. These connectome patterns and the extracted brain connectomic measures can be leveraged for potential clinical translational research such as the precision medicine (Fernandes et al., 2017; Hampel et al., 2019) linked to cognitive decline (Sorrentino, Rucco, Lardone, et al., 2021) and to Parkinson’s disease (Romano et al., 2022).

Brain fingerprinting shows great promise as a predictor of mental health outcomes and for that reason, it is explored under various neuroimaging modalities. Recently, few studies have started to explore connectome fingerprinting in different functional neuroimaging modalities, such as functional Near-Infrared Spectroscopy (fNIRS) (Rodrigues et al., 2019), electroencephalography (EEG) (Demuru and Fraschini, 2020), and magnetoencephalography (MEG) (Demuru et al., 2017; Sareen et al., 2021). It is important to underline here that brain fingerprinting research demands the access in a test-retest cohort (repeat scans) as a way to use the feature dataset derived from the first scan as a baseline database with the subject’s identity and the feature dataset extracted from the second scan for validation purposes of subject’s identification.

Another aim of my study was to investigate the repeatability of the Laplacian spectrum of SBN across alternative graph construction schemes. Complementary to the repeatability of Laplacian eigenvalues, I performed an identification analysis (brain fingerprinting) across pairs of scans where the second scan consists of the ‘target’ session and the first scan the ‘database’ session (Fig.4). Iteratively, one individual’s Laplacian eigenvalue was selected from the target set and compared against the N subject-specific Laplacian eigenvalues profile in the database set to find the Laplacian profile that was maximally similar. As a proper dissimilarity distance, I adopted the X^2^ statistics (Rubner, 2000). I followed a similar analysis on the Laplacian eigenvectors (harmonics) employing X^2^ statistics. The final outcome of this process is an identity matrix with 1s if the identity had been predicted correctly and 0s if it did not. Finally, I summed up the total number of corrected identifications per graph construction scheme and further divided by the total number of subjects to express the accuracy (performance) of the whole brain fingerprinting process. For a comparison purpose, I investigated the performance in terms of brain fingerprinting of the structural properties of SBN across alternative graph construction schemes separately for communities, 3,4-motifs, bipartiteness, and the total number of odd-cycle motifs and also in an ensemble way. I adopted proper metrics for every structural property such as normalized mutual information (MI) for the communities, the X^2^ statistics for the 3,4-motifs, the Euclidean distance (ED) for the bipartiteness and the X^2^ statistics for the total number of odd cycle motifs. For comparison purposes of previous studies and the extracted aforementioned features, I followed a brain connectome fingerprinting approach using Portrait Divergence metric as a proper graph distance metric applied over individual SBN (Bagrow and Bollt, 2019). We employed it in a previous systematic evaluation of fMRI data-processing pipelines for consistent functional connectomics (Luppi et al., 2024).

**Fig. 4.**
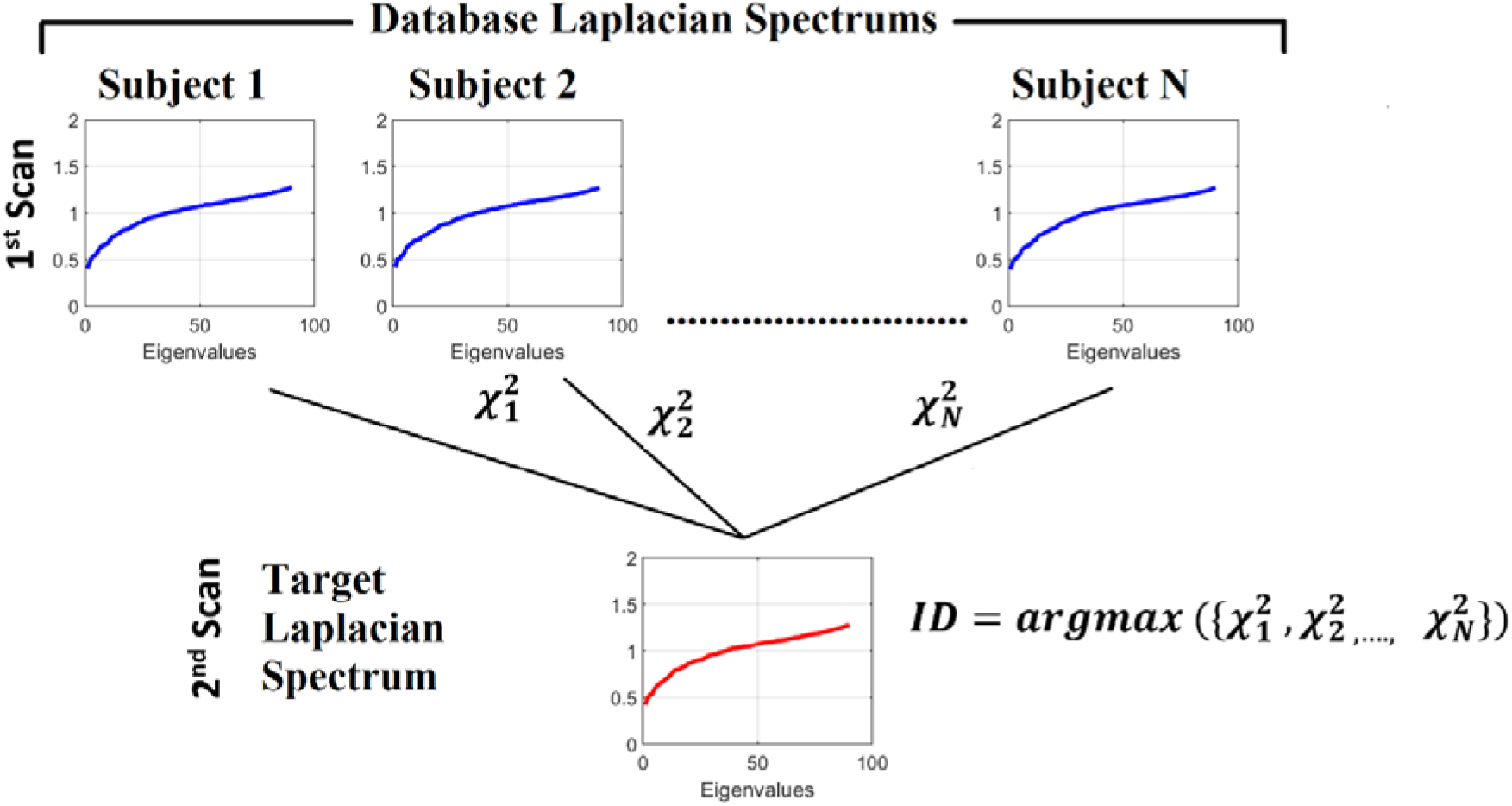
Identification analysis procedure. Identification procedure. Given a query Laplacian eigenvalue profile from the target set, I estimated the Chi-square histogram distance between this Laplacian eigenvalue profile and all the Laplacian eigenvalue profiles in the database. The predicted identity ID* is the one with the highest Chi-square histogram distance value (argmax) or the highest ED or the highest MI. In a similar way and with a proper statistical measure, I followed this identification approach for multiple structural properties.

### 2.6 Statistical Analysis

Below, I summarized the statistical analysis followed in my study.

#### Robustness of measurements with random rewired networks (surrogates)

To investigate the effect of adding small topological ‘noise’ to the structural brain networks to the Laplacian spectrum, I randomly rewired 5% of the edges while maintaining the degree and the strength of every node (Maslov and Sneppen, 2002). Ten thousand surrogate null network models were generated per subject, scan, and graph construction scheme (randmio_und_connected with iter = 10, Rubinov and Sporns,2010). All the properties estimated over original SBN and the relevant graph Laplacian spectrum were compared those obtained from the surrogate SBN and the related Laplacian spectrum.

#### Repeatability of Laplacian Eigenvalues

I employed Pcc between individual Laplacian eigenvalues between scans per graph construction scheme (within graph construction schemes) and Pcc between individual Laplacian eigenvalues across graph construction schemes in a pair-wise fashion (between graph construction schemes). Also, I applied a Wilcoxon Signed Rank-Sum test between the two sets of subject-specific Pcc values on the subject level that will support at which degree the Laplacian eigenvalues are highly dependent on the graph construction scheme.

#### Repeatability of Important nL-based properties

The repeatability of the Synchronizability and Laplacian energy per graph construction scheme and between the two scans was quantified with the absolute difference between the two scans. The original values were compared with the surrogate ones per graph construction scheme and scans. A p-value is assigned to the original values by a direct comparison with ten thousand surrogate values.

I applied a Wilcoxon Signed Rank-Sum test for both Synchronizability and Laplacian Energy properties between the two scans.

#### Laplacian Spectrum Properties Smaller Eigenvalues

I estimated group-mean λ_2_ (Fiedler value) and group-mean number of communities defined by the eigen-difference of Laplacian eigenvalues (Eigengap method) and by the k-Means clustering applied over the first eigenvectors. These group-means were averaged first across scans and then across subjects. Group-mean λ_2_ and the number of communities defined by the two methods were compared with the surrogate number of communities. To quantify the similarity of graph communities between the two scans (repeatability) per graph construction scheme and in both methods, I employed MI as a proper measure. The original MI values were also compared with surrogate MI values for both methods by adopting a Wilcoxon Rank-Sum test.

#### Medium Eigenvalues

I estimated the group-mean relative frequency (RF) linked to peak around one (λ = 1) across subjects for every graph construction scheme, at first averaged between scans. The original RF values were compared with the surrogate ones by adopting a Wilcoxon Rank-Sum test. As I aforementioned, I applied a multi-linear regression analysis between the RF and the total amount of each of the 3,4-motif and the Matching Index (MIN) across the SBN independently per scan. I estimated Pearson’s correlation coefficient between the nodal weighted clustering coefficient and the nodal 3-motifs distribution across graph construction schemes. Complementary, I estimated Pearson’s correlation coefficient in a pairwise fashion between nodal motif frequency of occurrence across the five graph construction schemes. I followed this approach independently per subject, scan and for each of the two 3-motifs and six 4-motifs. These Pcc correlations were averaged across scans first and afterwards across subjects. To compare my findings with those present in de Lange et al., (2016), I estimated the Pearson’s correlation coefficient between the MIN and the RF per graph construction scheme and scan and I presented the mean across graph construction scheme averaged across scans.

#### Largest Eigenvalues

In summary, I estimated the group-averaged of bipartivity index b_s_, of largest eigenvalue λn and of odd-cycles of various lengths across subjects for every graph construction scheme, at first averaged between scans. The original values of bs, of λn and of the exhaustive quantification of odd-cycles were compared with the surrogate ones by adopting a Wilcoxon Rank-Sum test. Complementary, I applied a multi-linear regression analysis between the largest eigenvalue λn and the bs plus the total number of odd-cycles of length 3, 5 and 7 across the SBN independently per scan.

#### Repeatability of Laplacian Eigenvectors

D^LS^ is estimated between scans and graph construction scheme. The original group-mean D^LS^ values for every graph construction scheme was compared with the surrogate ones.

#### Integrative, segregative, and degenerate harmonics of SBN

ELD is estimated between brain scans independently for the three regimes and graph construction scheme. I compared original ELD values with the surrogate ones independently for the three regimes and across graph construction schemes. I applied a Wilcoxon Rank-Sum test between the three regimes of harmonics in a pairwise fashion (3 pairs) and within every graph construction scheme and also per regime of harmonics across the graph construction schemes (5×4/2 = 10 pairs).

#### Multiple Comparison Correction

I applied the false discovery rate (q = 0.01) to correct for multiple comparisons.

## 3. Results

### 3.1 High Repeatability of Laplacian Eigenvalues

Laplacian eigenvalues were highly repeatable across the five graph construction schemes (within graph construction schemes;,). The between graph construction schemes correlation was with a. The Wilcoxon Signed Rank-Sum test revealed a strong difference in Pcc values derived from the within graph construction schemes with those Pcc values extracted from the between graph construction schemes (p-value = 0.0021). Fig.5 illustrates the Pcc values between all the combinations across the five graph construction schemes and scan sessions for subject 1. In the main diagonal, one can see the high Pcc values (repeatability level), and the relevant p-value between scans derived from the same graph construction scheme. The off-diagonal Pcc values refers to the between-session and graph construction schemes.

**Fig. 5.**
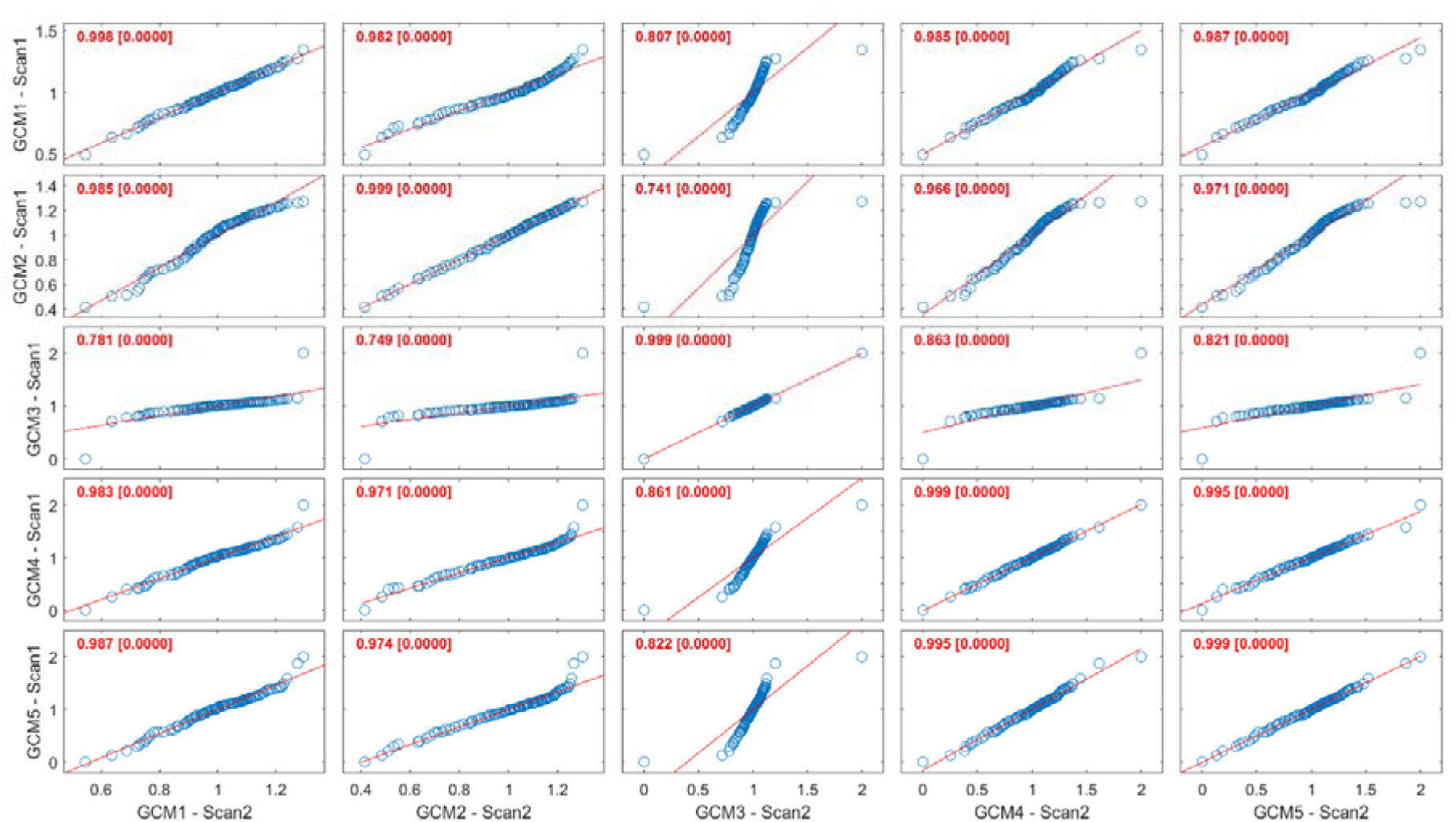
Pcc values between all the combinations across the graph construction schemes and scan sessions for subject 1. The main diagonal reports the repeatability estimations of the Laplacian eigenvalues (within-session and graph construction scheme) while the off-diagonal refers to the between-session and graph construction schemes.

### 3.2 Repeatability of the Laplacian Eigenvalue Properties

Fig.6 summarizes the group-mean and absolute between-scan differences of the Synchronizability and the Laplacian Energy for every graph construction scheme. Interestingly, the range of Synchronizability shows a higher dependency on the graph construction scheme compared to the Laplacian energy. My observations are supported by a direct comparison of original values with the surrogate-based Laplacian properties. P-values for both Synchronizability and Normalized Laplacian Energy were significant compared to surrogates across graph construction schemes (p < 0.001). The smallest group-mean between-scan difference for Synchronizability is shown for the 9m-OMST (p-value = 0.0041) and NS-OMST (p-value = 0.0021) graph construction schemes and the Laplacian energy is shown for 9m-OMST (p-value = 0.0032) and NS – OMST (p-value = 0.0022) graph construction schemes.

**Fig. 6.**
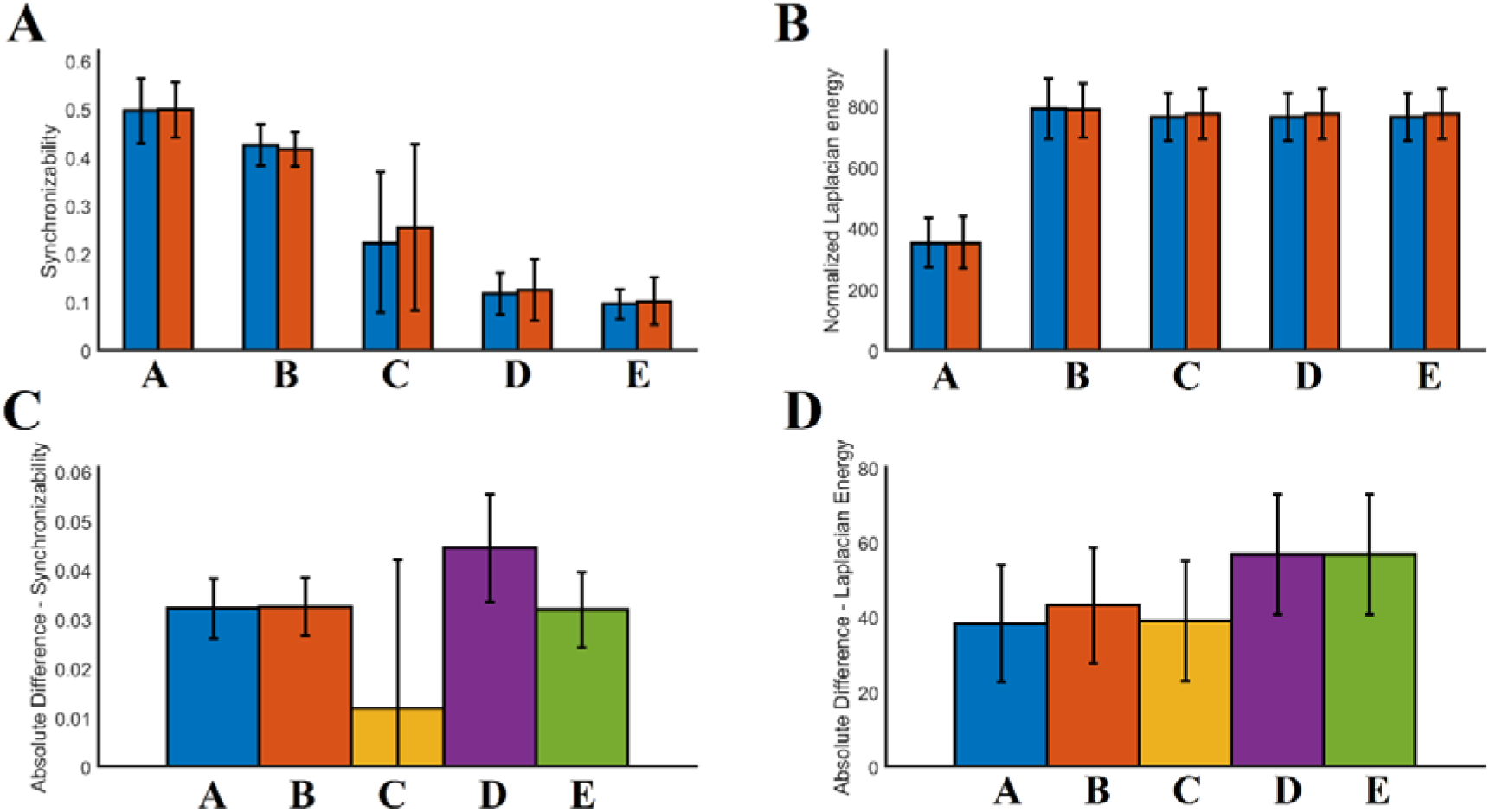
Illustration of the group-mean Synchronizability and Laplacian Energy for every graph construction scheme. A. Group-mean Synchronizability, B. group-mean normalized Laplacian Energy, C. Group-mean between-scan absolute difference of Synchronizability, and D Group-mean between-scan absolute difference of Laplacian Energy. Letters from A to E refer to the five graph construction schemes defined in Table 2. In A and B, blue/red colors refer to the first and second scan sessions, correspondingly.

### 3.3 Laplacian Sub-spectrum findings Smaller Eigenvalues

Table 3 summarizes the group-mean λ_2_, the group-mean number of modules estimated with the method of eigen-difference of Laplacian eigenvalues (Eigengap method) and by the K-Means clustering applied over the first eigenvectors. For the first two graph construction schemes (NS-OMST, 9m-OMST) compared to the surrogate null models, the average λ_2_ revealed that the original SBN can be subdivided in two subnetworks, while the eigengap and the K-Means approaches for the detection of communities showed significant different findings compared to the surrogates (p < 0.001).

Table 4 tabulates the MI of between-scan communities affiliations extracted with both methods and in every graph construction scheme. For the first two graph construction schemes (NS-OMST, 9m-OMST), the MI values for the K-Means algorithm are high, while the statistical comparison of the MI between the K-Means and the eigengap algorithms (p < 0.001) untangled the K-Means algorithm as a better approach compared to the eigengap. The communities extracted with K-Means applied over the SBN constructed with the 9m-OMST method showed a high similarity with our previous study (*MI* = 0.91 ± 0.04) where numerous graph partition algorithms were applied on the same set (Dimitriadis et al., 2021).

**Table 4.**
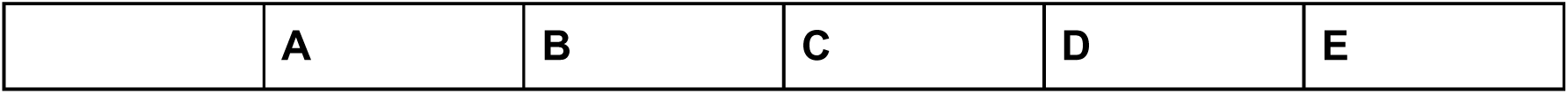
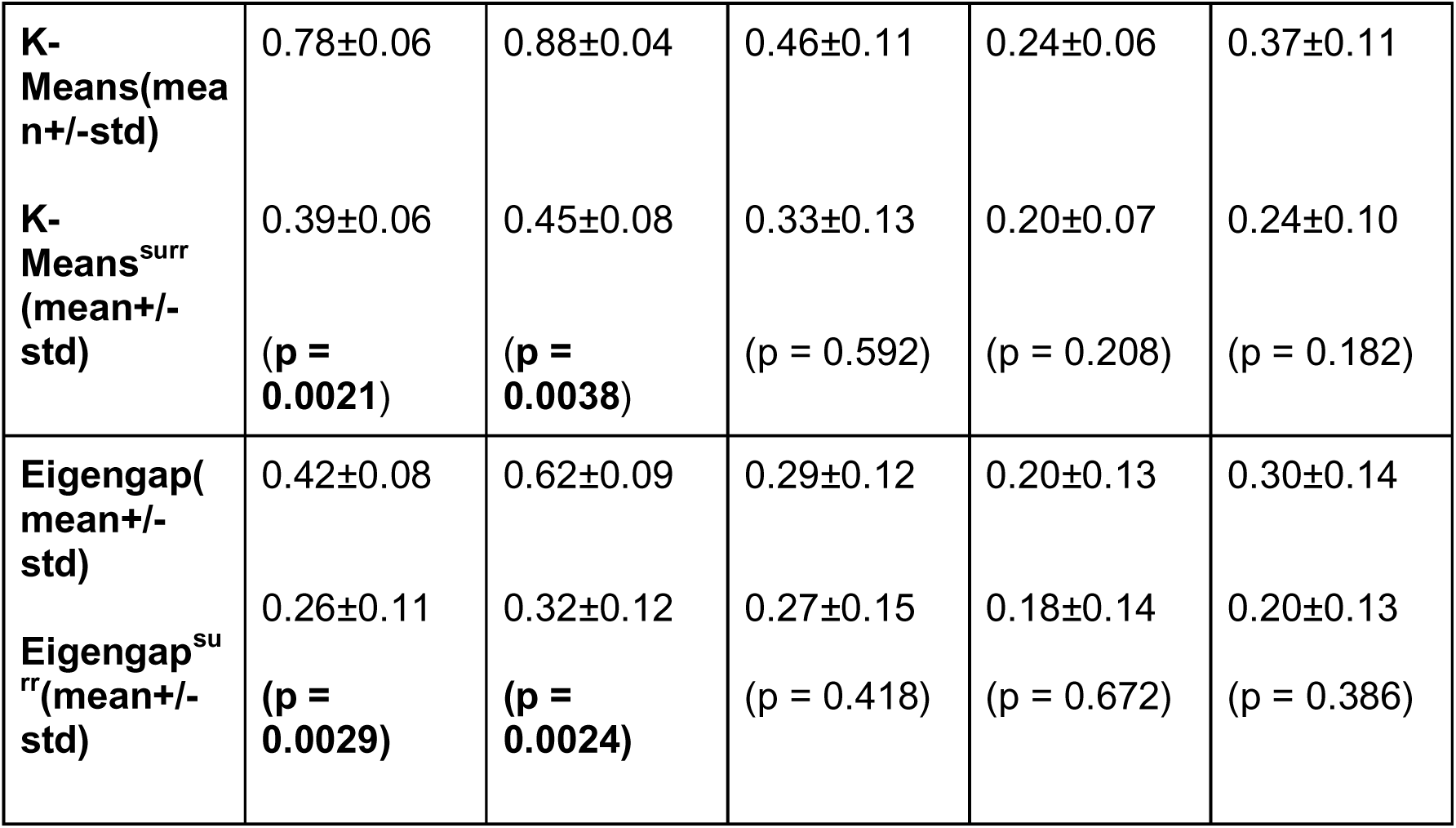
The between-scan MI of the communities extracted with eigen-difference and K-Means clustering per graph construction schemes. I underlined with bold, the p-values that showed significant differences compared to the surrogate-based p-values. (Letters from A to E refer to the five graph construction schemes defined in Table 2.

#### Medium Eigenvalues

The Laplacian spectrum of the graph construction schemes showed a single clear smooth peak as presented in Fig.2. This peak around 1 suggests a number of motif duplications where its height is relevant to this number. The relationship between the relative frequency (RF) linked to peak at λ = 1 and the total number of unique 3 and 4 motifs and the Matching Index (MIN) is described below. The Laplacian spectrum of structural brain networks across subjects, scans, and graph construction schemes didn’t show any other clear peaks indicative of recurrent addition of motifs. My findings are supported by the surrogate analysis where peaks of significant lower amplitude were observed in surrogated Laplacian spectrums. All distributions showed a peak around 1 while the group-mean relative frequency (RF) related to this peak didn’t show differences across methods (p > 0.05, Bonferroni correction) but showed difference between original and surrogate null models (Table 5).

**Table 5.** Group-mean relative frequency (RF) linked to peak at λ = 1 across subjects for every graph construction scheme. I underlined with bold, the p-values that showed significant differences compared to the surrogate-based p-values. (Letters from A to E refer to the five graph construction schemes defined in Table 2.

|  | A | B | C | D | E |
| --- | --- | --- | --- | --- | --- |
| **RF(mean+/-std)** | 2.36±0.05 | 2.67±0.06 | 2.59±0.13 | 2.54±0.12 | 2.42±0.15 |
| **$RF^{surr}$ (mean+/-std)** | 1.29±0.12 | 1.65±0.08 | 1.65±0.14 | 1.63±0.15 | 1.55±0.14 |
|  | (p = **0.32x10<sup>-12</sup>**) | (p = **0.83x10<sup>-11</sup>**) | (p = **0.64x10<sup>-10</sup>**) | (p = **0.89x10<sup>-10</sup>**) | (p = **0.42x10<sup>-12</sup>**) |

The multi-linear regression analysis revealed a significant trend between the RF and the motifs only for the 9m-OMST graph construction scheme. The following equation described the relationship between the RF and the two **3-motifs, the first four 4-motifs and the Matching Index (MIN)** (see Fig.3A):

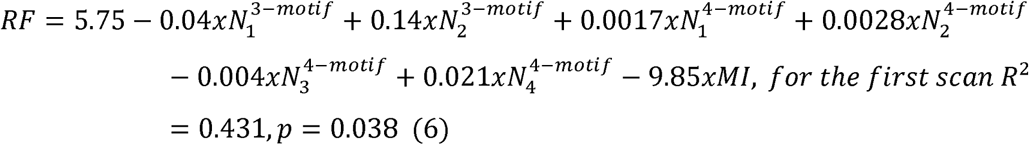

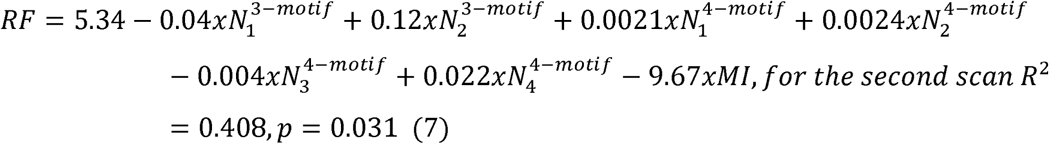

Mean and standard deviations of motifs and MI for each scan (first / second scan).

First scan :

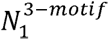 = 72.43 ± 17.31,

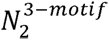 = 15.43 ± 4.72,

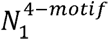 = 492.70 ± 175.23,

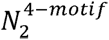 = 129.78 ± 48.36,

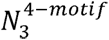 = 231.08 ± 93.58,

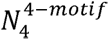 = 10.49 ± 4.51,

*MI* = 0.24 ± 0.0086

Second scan:

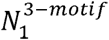 = 70.81 ± 15.33,

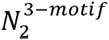 = 15.45 ± 4.26,

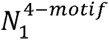 = 470.36 ± 148.35,

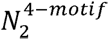 = 123.76 ± 42.13,

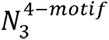 = 227.01 ± 82.63,

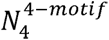 = 9.69 ± 3.60,

*MI* = 0.24 ± 0.0078

As per motifs, I encountered the total amount of either 3 or 4-motifs detected across the individual structural connectome per scan. Typically, this is the sum of distribution showed in Fig.7,8 in columns across graph construction schemes. Figs 7-8 show the averaged across subjects and scans motif fingerprint of every node (ROI) across the five graph construction schemes for each of alternative 3,4-motifs.

**Fig. 7.**
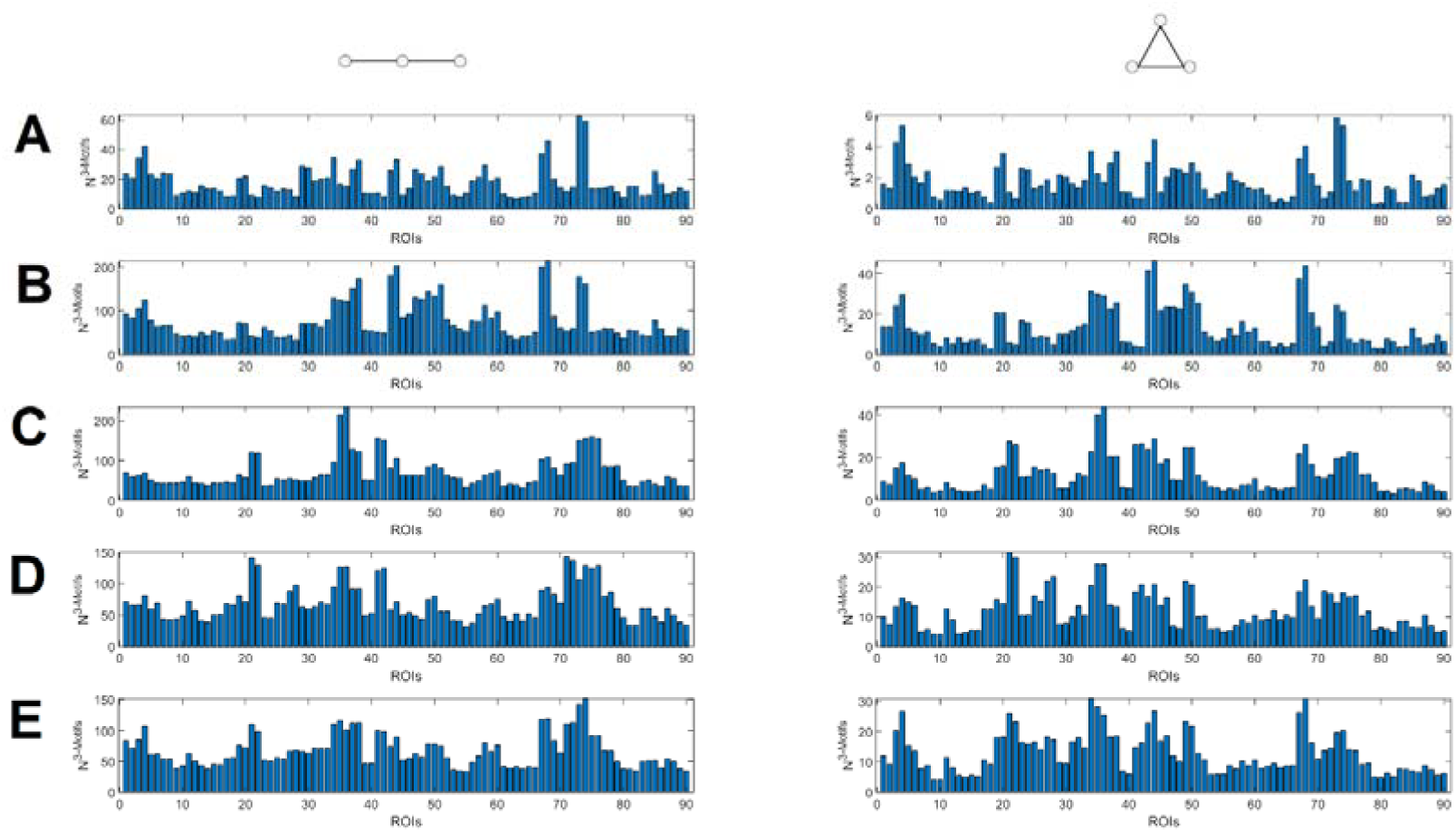
Group and scan averaged motif fingerprint of every ROI across the five graph construction schemes (A-E) and the two 3,motifs.

**Fig. 8.**
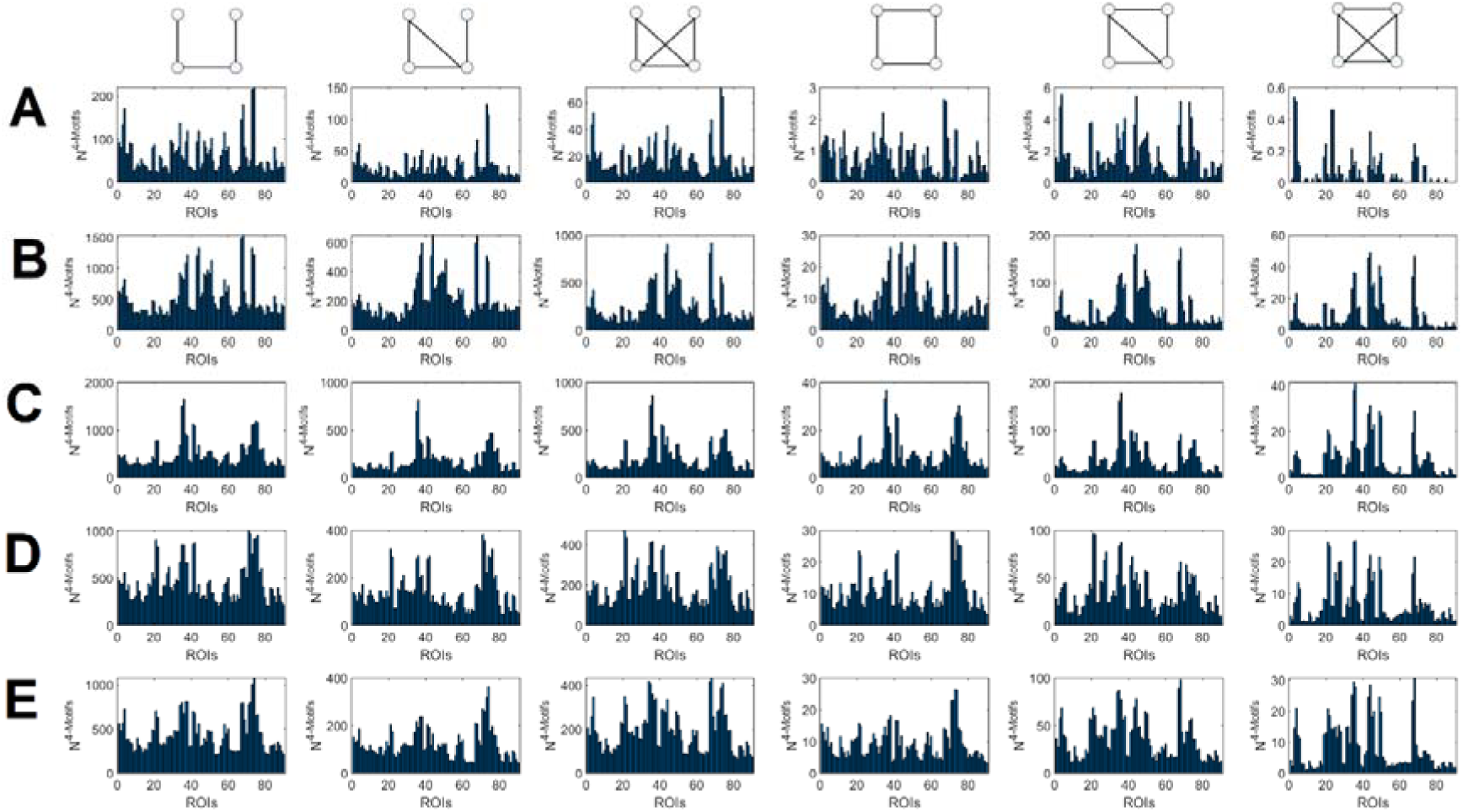
Group and scan averaged motif fingerprint of every ROI across the five graph construction schemes (A-E) and the two 4,motifs.

The correlation of the MIN with the RF was 0.54 ± 0.12 averaged across the graph construction scheme. This finding underlines how network symmetry quantified with MIN shapes the central peak of the graph Laplacian spectrum.

Fig.9 illustrates the group-averaged Pearson’s correlation coefficient between the nodal weighted clustering coefficient and the nodal 3-motifs distribution (Fig.7) across the five graph construction schemes. I first estimated the mean of correlations between scans per subject across the graph construction schemes. A positive correlation between the second 3-motif and weighted clustering coefficient was consistently observed across graph construction schemes while a mixed sign of correlation was detected for the first 3-motif (Fig.9).

**Fig. 9.**
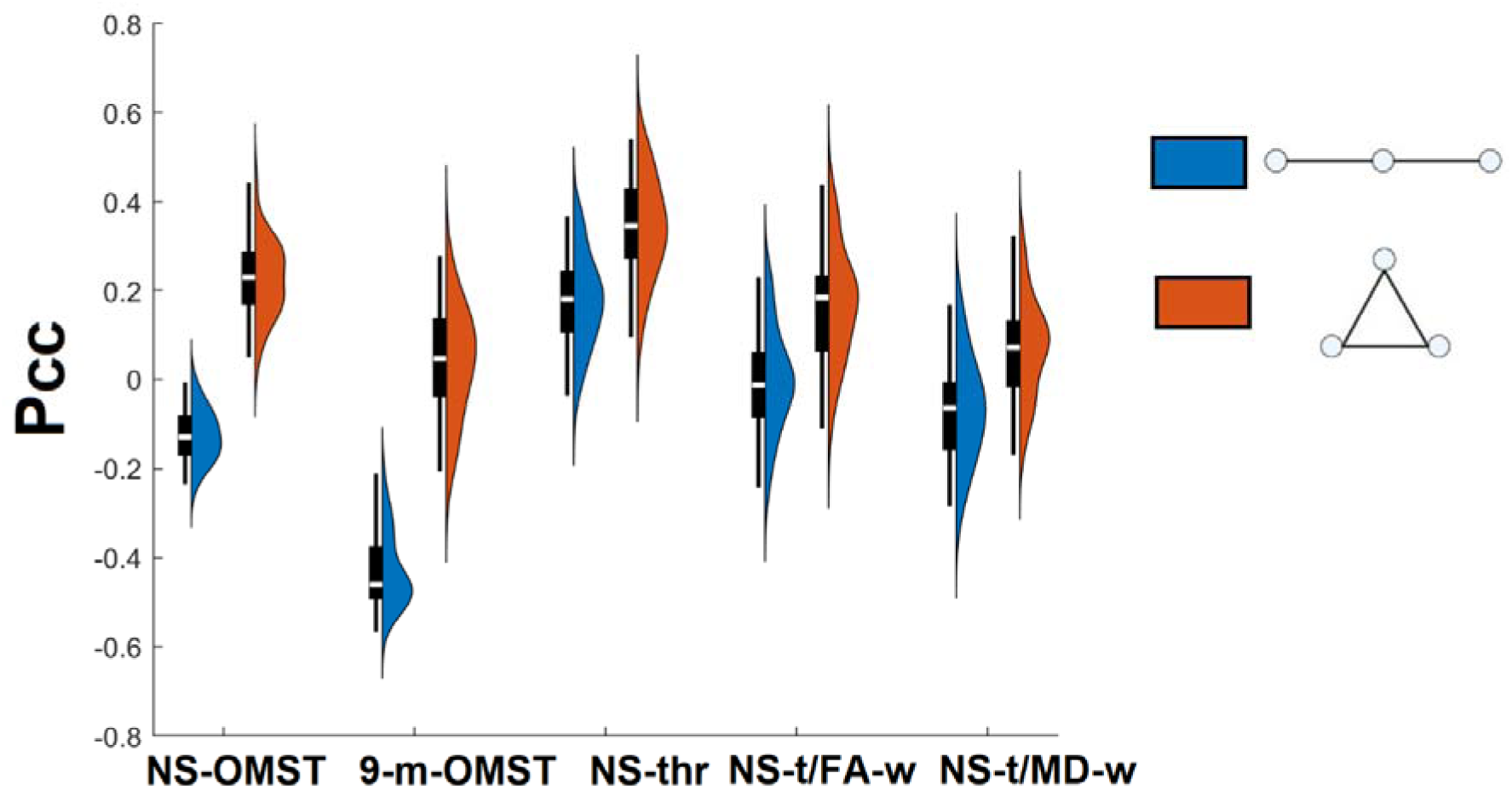
Group-averaged Pearson’s correlation coefficient between the nodal weighted clustering coefficient and the two nodal 3-motifs distribution as shown in Fig.7 Mean values and standard deviation refer to the group level (Pcc – Pearson’s Correlation Coefficient).

Fig.10 illustrates the group-averaged Pearson’s correlation coefficient between the motif frequency of occurrences across the graph constructions schemes independently for each of the two 3-motifs (Fig.7). Similarly, Fig.11 illustrates the group-averaged Pearson’s correlation coefficient between the motif frequency of occurrences across the graph constructions schemes independently for each of the six 4-motifs (Fig.7). I first estimated the mean of correlations between scans per subject across graph construction schemes. In both 3,4-motifs, a strong positive correlation was consistently observed between the 9-m-OMST, NS-thr and NS-t/MD-w. This means that the individual SBN shares a large number of common local topologies that is reflected to the global network level.

**Fig. 10.**
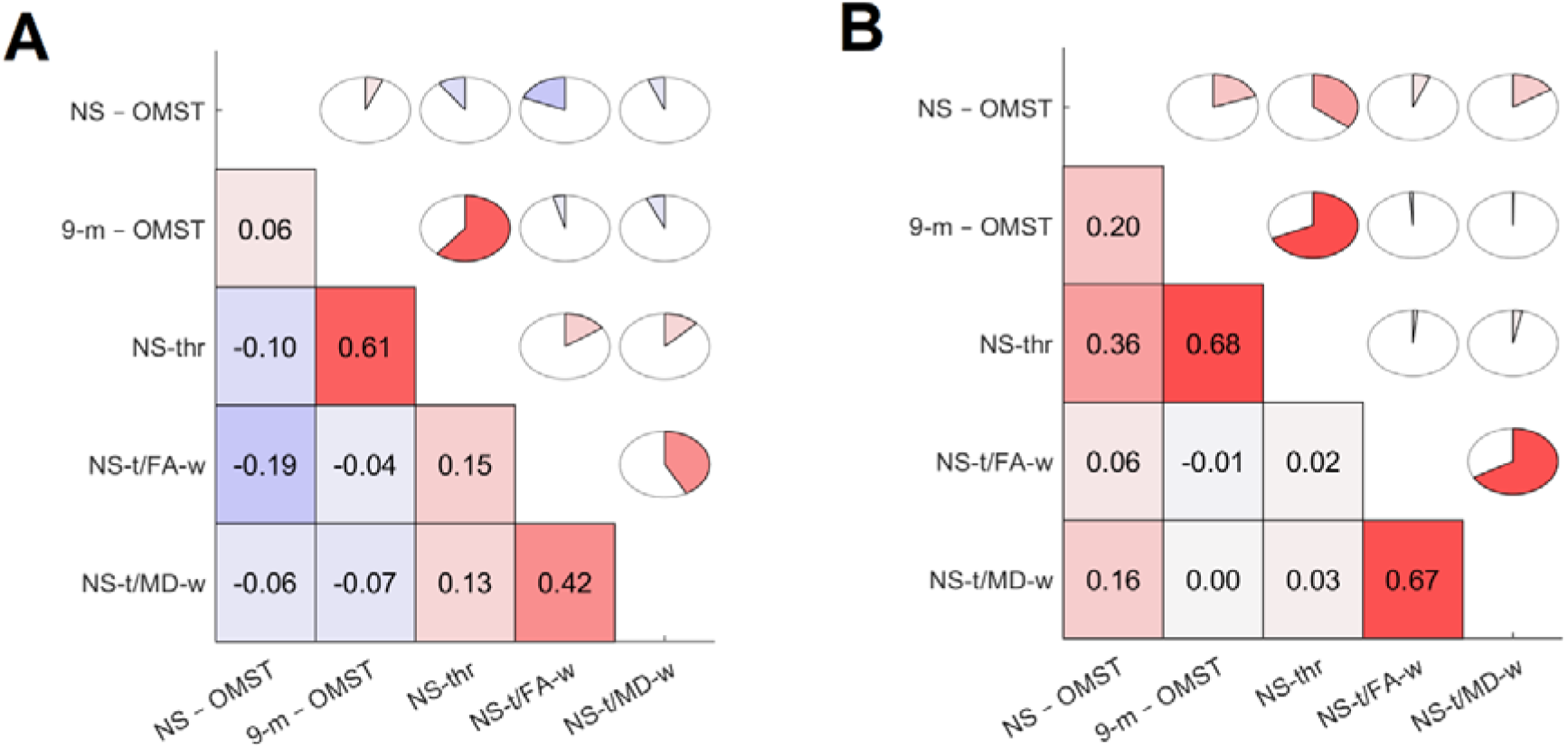
Group-averaged Pearson’s correlation coefficients of the motif frequency of occurrence between pairs of graph construction schemes for each of the two 3-motifs. A and B refer to the two 3-motifs as demonstrated in Fig.7.

**Fig. 11.**
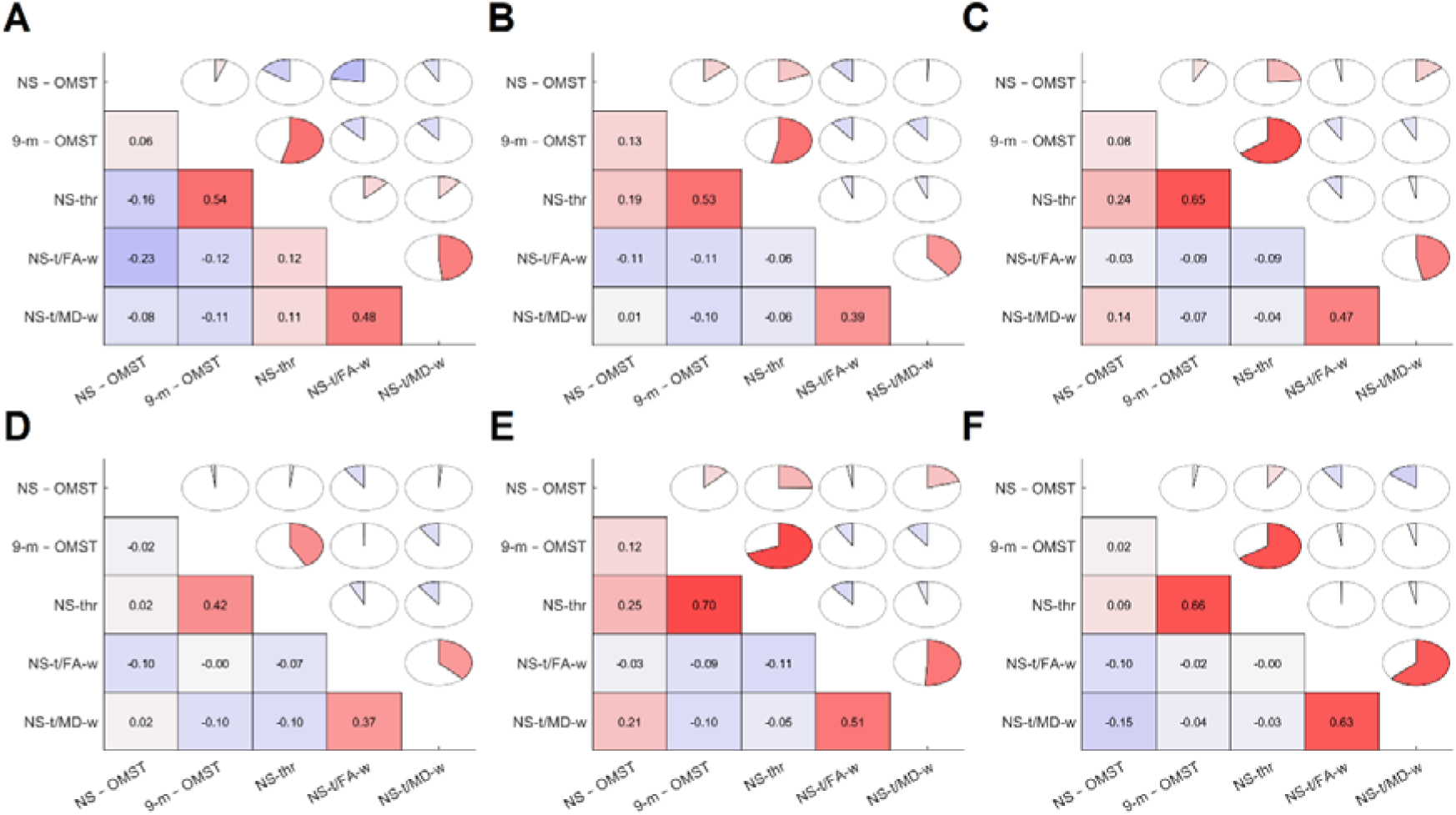
Group-averaged Pearson’s correlation coefficients of the motif frequency of occurrence between pairs of graph construction schemes for each of the six 4-motifs. A – F refer to the six 4-motifs as demonstrated in Fig.8.

#### Largest eigenvalues

Table 6 summarizes the group-mean largest eigenvalue λ_n_ across graph construction schemes which differs significantly from the largest eigenvalue relevant to the random networks. The largest eigenvalue of the Laplacian spectrum informs us of the level of ‘bipartiteness’ of the most bipartite subpart of the network, which is closely related to the number of odd cyclic motifs in the network. Visual inspection of the associated eigenvector linked to the largest eigenvalue across the cohort, scans, and in the first two graph construction schemes (NS-OMST, 9m-OMST) with respect to communities as detected in (Dimitriadis et al., 2021; see Fig.5), is highly localized in modules 8 and 9.

**Table 6.** Group-mean large eigenvalue λ_n_. (Letters from A to E refer to the five graph construction schemes defined in Table 2). **I underlined with bold, the p-values that showed significant differences compared to the surrogate-based p-values** (Letters from A to E refer to the five graph construction schemes defined in Table 2)

|  |  |  |  |  |  |
| --- | --- | --- | --- | --- | --- |
|  | **A** | **B** | **C** | **D** | **E** |
| $\lambda_n(\text{mean} \pm \text{std})$ | 1.32±0.05 | 1.33±0.02 | 1.31±0.02 | 1.29±0.03 | 1.28±0.02 |
| $\lambda_n^{\text{surr}}(\text{mean} \pm \text{std})$ | 1.09±0.06 | 1.08±0.03 | 1.07±0.04 | 1.05±0.05 | 1.03±0.05 |
|  | (p = **0.53x10<sup>-13</sup>**) | (p = **0.32x10<sup>-15</sup>**) | (p = **0.37x10<sup>-15</sup>**) | (p = **0.43x10<sup>-13</sup>**) | (p = **0.62x10<sup>-14</sup>**) |

Table 7-9 shows the group-mean odd-cycles (odd-cycles of length 3,5 and 7) fingerprint related to the total number of odd-cycles across the graph construction schemes. The odd-cycle fingerprint was significantly different for all the graph construction schemes compared to the random networks for the three lengths.

**Table 7.** Group-mean total number of the odd-cycles of length (*l*) = 3. (Letters from A to E refer to the five graph construction schemes defined in Table 2). **I underlined with bold, the p-values that showed significant differences compared to the surrogate-based p-values** (Letters from A to E refer to the five graph construction schemes defined in Table 2).

|  | A | B | C | D | E |
| --- | --- | --- | --- | --- | --- |
| *odd – cycles ($l = 3$)*<br>(mean±-std) | 53.24±2.94 | 463.37±117.69 | 609.56±101.79 | 77.82±12.51 | 69.51±13.78 |
| *odd – cycles ($l = 3$)<sub>surr</sub>*(mean±-std) | 31.51±2.37 | 294.72±75.45 | 472.83±92.45 | 33.77±13.62 | 40.92±8.62 |
|  | (p = **0.24x10<sup>-12</sup>**) | (p = **0.51x10<sup>-13</sup>**) | (p = **0.18x10<sup>-12</sup>**) | (p = **0.12x10<sup>-13</sup>**) | (p = **0.23x10<sup>-15</sup>**) |

**Table 8.**
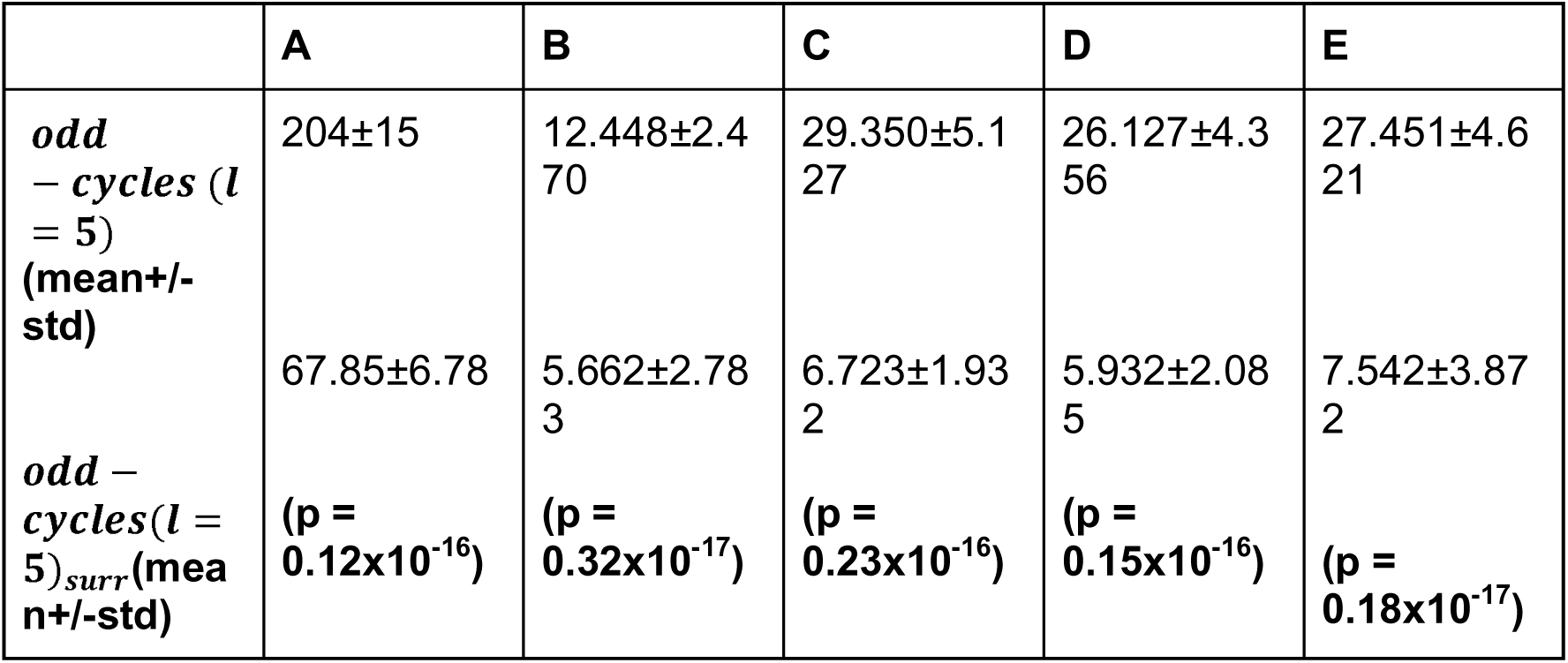
Group-mean total number of the odd-cycles of length (*l*) = 5. (Letters from A to E refer to the five graph construction schemes defined in Table 2). **I underlined with bold, the p-values that showed significant differences compared to the surrogate-based p-values** (Letters from A to E refer to the five graph construction schemes defined in Table 2).

|  | A | B | C | D | E |
| --- | --- | --- | --- | --- | --- |
| *odd*<br>– *cycles* (*l*<br>= 5)<br>(mean+/-<br>std) | 204±15 | 12.448±2.4<br>70 | 29.350±5.1<br>27 | 26.127±4.3<br>56 | 27.451±4.6<br>21 |
| *odd* –<br>*cycles* (*l* =<br>5)<sub>surr</sub>(mea<br>n+/-std) | 67.85±6.78<br>(p =<br>**0.12x10<sup>-16</sup>**) | 5.662±2.78<br>3<br>(p =<br>**0.32x10<sup>-17</sup>**) | 6.723±1.93<br>2<br>(p =<br>**0.23x10<sup>-16</sup>**) | 5.932±2.08<br>5<br>(p =<br>**0.15x10<sup>-16</sup>**) | 7.542±3.87<br>2<br>(p =<br>**0.18x10<sup>-17</sup>**) |

**Table 9.** Group-mean total number of the odd-cycles of length (*l*) = 7. (Letters from A to E refer to the five graph construction schemes defined in Table 2). **I underlined with bold, the p-values that showed significant differences compared to the surrogate-based p-values** (Letters from A to E refer to the five graph construction schemes defined in Table 2).

|  | A | B | C | D | E |
| --- | --- | --- | --- | --- | --- |
| *odd*<br>– *cycles* (*l*<br>= 7)<br>(mean+/-<br>std) | 798.10±88.<br>20 | 414.980 ±<br>337.180 | 1.765.000±<br>907.230 | 1.326.000±<br>675.341 | 1.452.000±<br>745.432 |
| *odd* –<br>*cycles* (*l* =<br>7)<sub>surr</sub>(mea<br>n+/-std) | 417.56±<br>59.14<br>(p =<br>**0.09x10<sup>-17</sup>**) | 217.354±<br>168.931<br>(p =<br>**0.21x10<sup>-17</sup>**) | 889.451±<br>392.753<br>(p =<br>**0.11x10<sup>-17</sup>**) | 712.562±<br>294.431<br>(p =<br>**0.12x10<sup>-17</sup>**) | 798.542±<br>334.231<br>(p =<br>**0.11x10<sup>-16</sup>**) |

Table 10 tabulates the group-mean bipartivity index b_s_ across the graph construction schemes. The bipartivity index b_s_ was significantly different only for the first two graph construction schemes (NS-OMST, 9m-OMST) compared to the random networks. Interestingly, the first graph construction scheme (NS-OMST) produced the highest bipartivity compared to the rest of graph construction schemes. Multi-linear regression analysis between the λn and the bs plus the three exhaustive estimation of odd-cycles of length = 3,5,7 across the graph construction schemes revealed interesting findings only for the first two graph construction schemes (Table 11). The multi-linear regression model for the 9m-OMST showed the highest R^2^ and the lowest p-value compared to the NS – OMST uncovering a relationship between RF and structural network properties expressed with bs and odd-cycles. Findings were consistent in both scans.

**Table 10.** Group-mean bipartivity index b_s_. (Letters from A to E refer to the five graph construction schemes defined in Table 2). **We underlined with bold, the p-values that showed significant differences compared to the surrogate-based p-values** (Letters from A to E refer to the five graph construction schemes defined in Table 2).

|  | A | B | C | D | E |
| --- | --- | --- | --- | --- | --- |
| **$b_s$(mean+/-std)** | 0.87±0.02 | 0.19±0.04 | 0.16±0.06 | 0.17±0.05 | 0.19±0.06 |
| **$b_s^{surr}$(mean+/-std)** | 0.15±0.05 | 0.06±0.05 | 0.09±0.04 | 0.10±0.04 | 0.10±0.07 |
|  | **(p = $0.46 \times 10^{-14}$)** | **(p = $0.56 \times 10^{-15}$)** | **(p = 0.732)** | **(p = 0.672)** | **(p = 0.707)** |

**Table 11.** Outcome of multi-linear regression analysis between the λn and the bs with the three odd-cycles (OC) of length (*l*) = 3,5,7. Every row corresponds to each scan (with ‘x’, I denote the arguments that didn’t overcome the statistical threshold of p-value < 0.05)

| | Intercept | $b_s$ | $OC^{l=3}$ | $OC^{l=5}$ | $OC^{l=7}$ | $R^2$ | p-value |
| --- | --- | --- | --- | --- | --- | --- | --- |
| **A** | 1.86 | x | - | 0.00090 | - | 0.312 | 0.0152 |
|  | 1.85 | x | 0.0042<br>-.0039 | 0.00085 | 0.000087<br>-<br>0.000077 | 0.295 | 0.0245 |
| **B** | **1.81**<br>**1.82** | **-0.057**<br>**-0.061** | **0.0012**<br>**0.0013** | **0.00025**<br>**0.00027** | -<br>**0.000017**<br>-<br>**0.000016** | **0.844**<br>**0.812** | **1.71e-12**<br>**2.65e-11** |
| **C** | 2.10<br>2.05 | x<br>x | x<br>x | x<br>x | x<br>x | 0.222<br>0.213 | 0.0819<br>0.0912 |
| **D** | 2.01<br>1.92 | x<br>x | x<br>x | x<br>x | x<br>x | 0.204<br>0.187 | 0.0721<br>0.0834 |
| **E** | 1.95<br>1.86 | x<br>x | x<br>x | x<br>x | x<br>x | 0.192<br>0.167 | 0.0654<br>0.0782 |

### 3.4 Repeatability of Laplacian Eigenvectors

Table 12 reports the group-mean between-scan **D^LS^** of the Laplacian eigenvectors for every graph-construction scheme. 9m-OMST graph construction scheme showed the smallest group-mean D^LS^ followed by the FA-t/NS-w but without reaching the significant level (see Fig.12; p-value < 0.05, Bonferroni corrected). My findings are supported also by the direct comparison of original D^LS^ values with the surrogate D^LS^ values (p-value = 0.0057 & p-value = 0.0043, for 9m-OMST and NS-OMST, correspondingly). An overview of the two Laplacian eigenvectors (connectomic harmonics) from both scan sessions for 9m-OMST are illustrated in Fig.13.

**Fig. 12.**
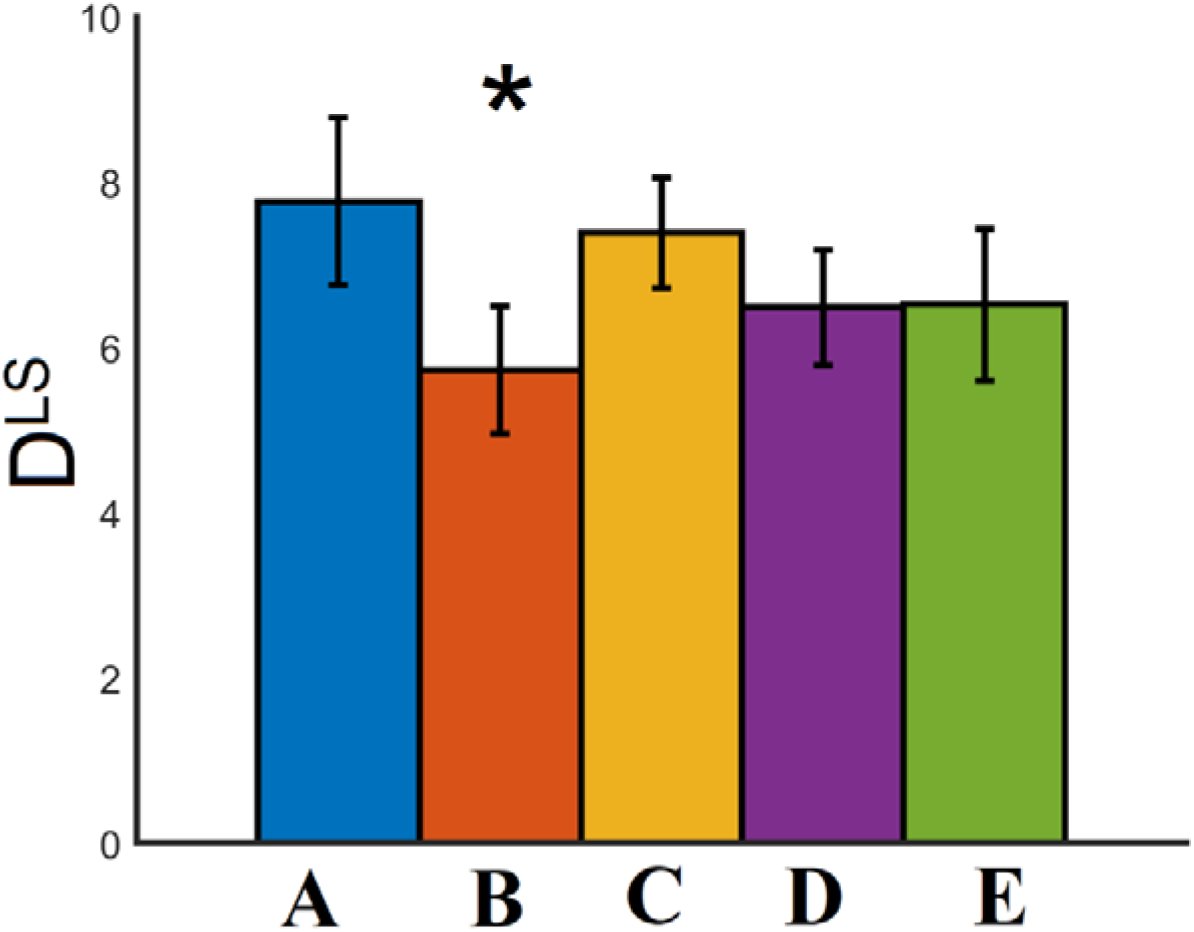
Illustration of the group-mean D^LS^ for every graph construction scheme. (* denotes the statistical difference of D^LS^ for the 9m-OMST method versus the four methods).

**Fig. 13.**
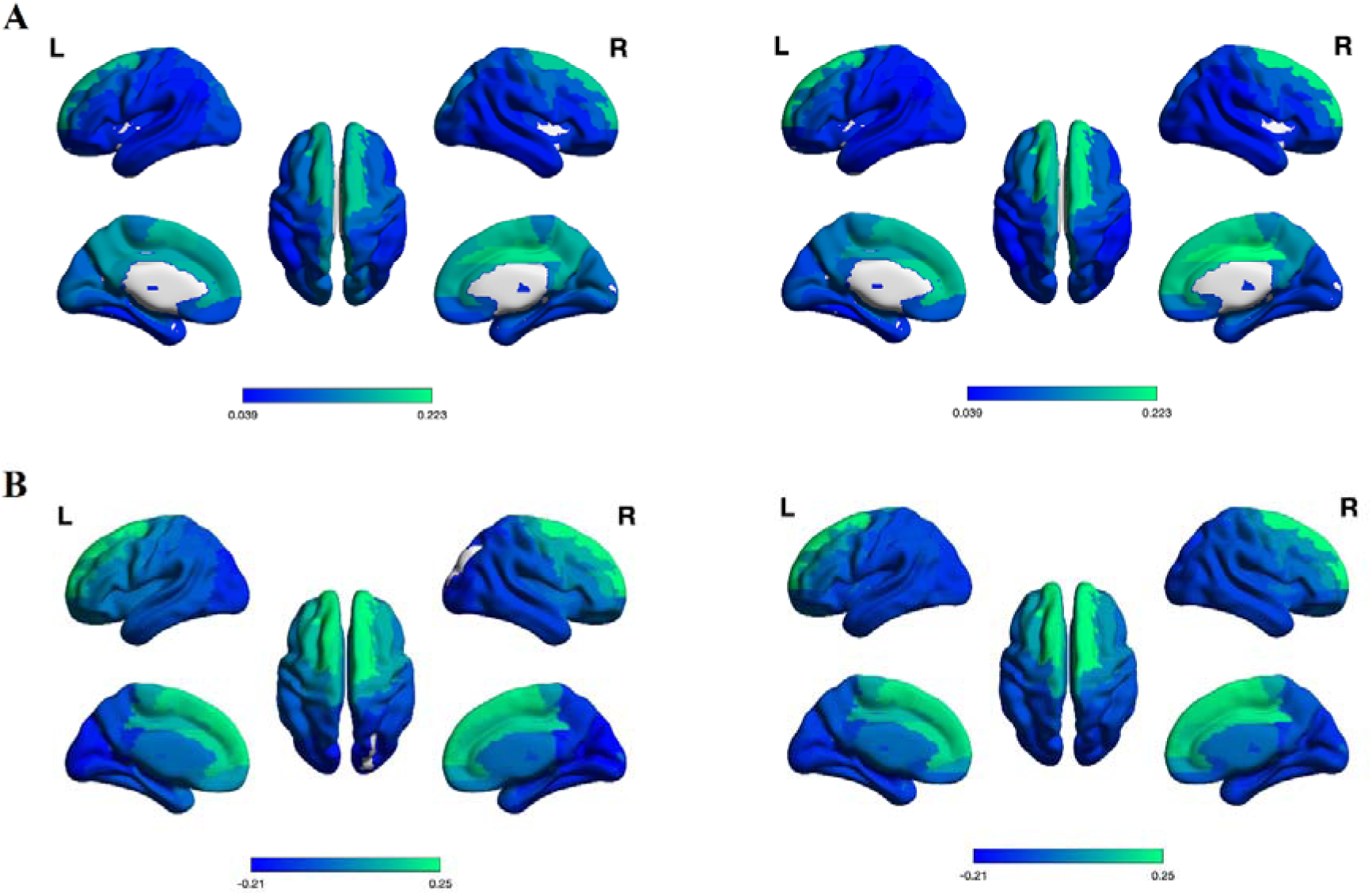
Overview of the two Laplacian eigenvectors (connectomic harmonics) from both scan sessions for 9m-OMST method extracted from subject 1. A. First eigenvector from scan 1 (left) and scan 2 (right) B. Second eigenvector from scan 1 (left) and scan 2 (right)

**Table 12.**
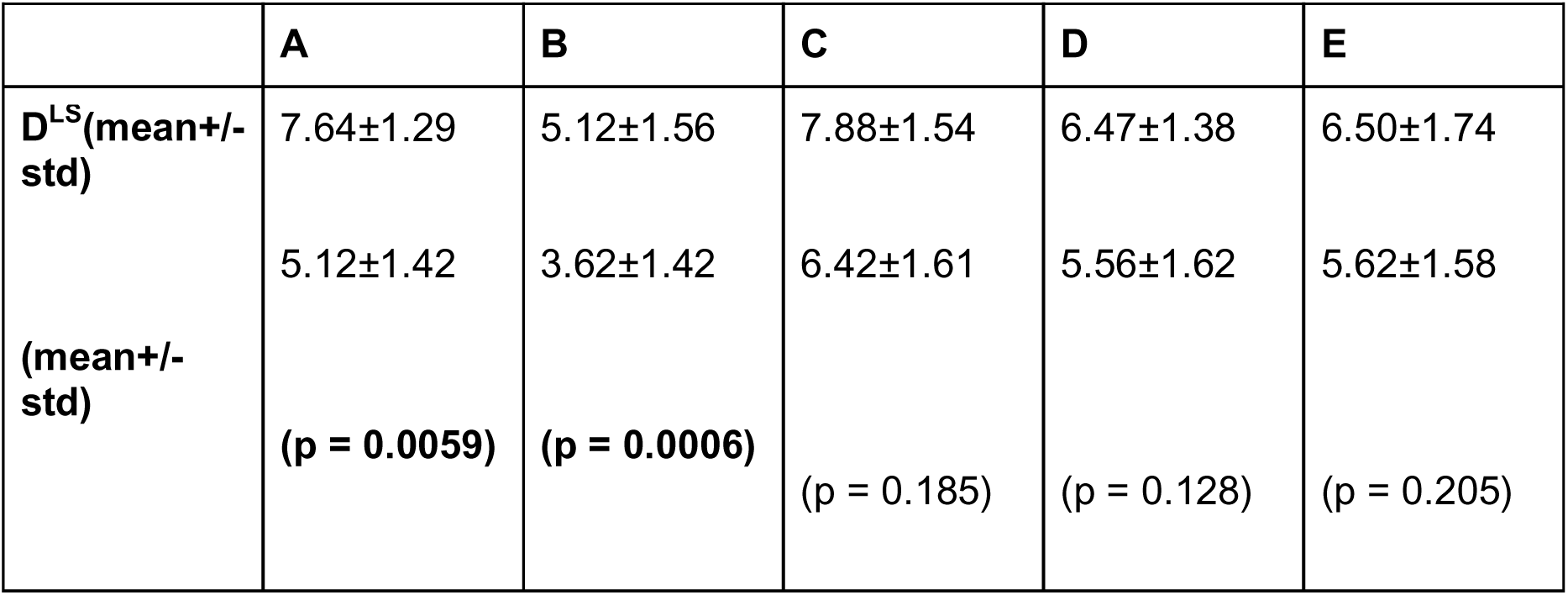
Group-mean between scan D^LS^ of the Laplacian eigenvectors for every graph construction scheme. I underlined with bold, the p-values that showed significant differences compared to the surrogate-based p-values. (Letters from A to E refer to the five graph construction schemes defined in Table 2).

### 3.5 Repeatability of the Integrative, segregative, and degenerate harmonics of SBN using ELD

Fig.14 illustrates the group-averaged ELD values for every regime of harmonics across the graph construction schemes. It is clear that the repeatability of harmonics’ partitions is higher for the integrative harmonic, middle for the degenerate and lower for the segregative. Group-averaged ELD values for every regime of harmonics and across graph construction schemes were significantly different compared to the surrogate ELD values (p < 0.01 x 10^-5^, Bonferroni corrected). Statistical analysis between the three regimes of harmonics in a pairwise fashion and within every graph construction scheme showed significant differences for every pair and graph construction scheme (p < 0.01 x 10^-12^, Bonferroni corrected). The comparison of ELD values per regime of harmonics across the graph construction schemes revealed interesting trends. No significant difference was detected across the graph construction schemes for integrative and degenerate harmonics while group-averaged ELD was statistically lower for segregative harmonics for 9-m-OMST graph construction scheme compared to the rest (p < 0.02 x 10^-12^, Bonferroni corrected).

**Fig. 14.**
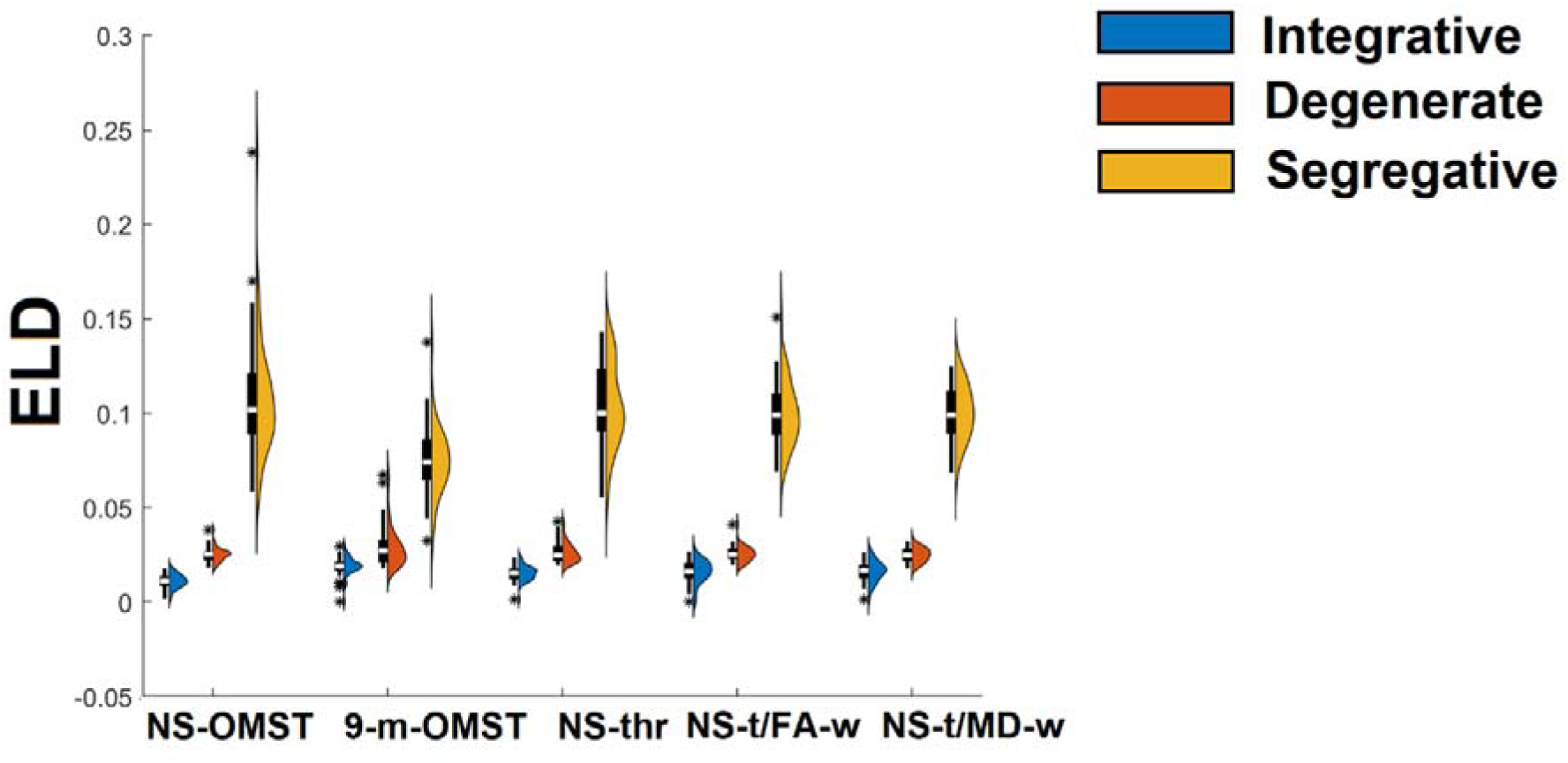
Repeatability of harmonics’ partitions. Group-averaged ELD values for every regime of harmonics across the graph construction schemes.

### 3.6 Brain Fingerprinting

I presented the various inputs in the brain fingerprinting approach underling in parenthesis the adopted measure. My analysis succeeded in an accurate identification of subjects id (100%) on the test set (second session) on every graph construction scheme employing the Laplacian spectrum (Laplacian eigenvalues) as a feature vector (X^2^ statistics). However, the identification accuracy employing Laplacian eigenvectors (harmonics) showed a 100% level only for the 9m-OMST.

Similarly, I succeeded an absolute identification of subjects id (100%) across every graph construction scheme using the whole structural connectome as a 2D tensor using Portrait Divergence (PID) as a proper graph theoretic comparison metric. Complementary, Table 13 shows the Identification accuracy of structural properties of SBN across alternative graph construction schemes. Across the five studying structural properties, the highest accuracies were detected for communities (MI), and 3,4-motifs (X^2^ statistics) across the five construction schemes while the highest performance was detected for 9m-OMST (B), and NS-OMST (A) with the former to get higher accuracies. The identification accuracy for Bipartiveness (ED) was too low while and for the odd-cycle motifs (X^2^ statistics) were high only for the 9m-OMST (B), and NS-OMST (A). The combination of the outcome of brain identification strategy for communities, 3,4-motifs and odd-cycles of **length (*l*) = 3,5,7** (ensemble way) gave an absolute accuracy (100%) for the 9m-OMST (B), and a 94.59 for the NS-OMST (A).

**Table 13.** Identification accuracy of structural properties of SBN across alternative graph construction schemes. (Letters from A to E refer to the five graph construction schemes defined in Table 2).

|  | **A** | **B** | **C** | **D** | **E** |
| --- | --- | --- | --- | --- | --- |
| **Laplacian Eigenvalues ($X^2$)** | 100 | **100** | 100 | 100 | 100 |
| **Laplacian Eigenvectors ($X^2$)** | 72.97 | **86.49** | 59.46 | 56.76 | 56.76 |
| **SBN (PID)** | 100 | **100** | 100 | 100 | 100 |
| **Communities (MI)** | 83.78 | **89.19** | 56.76 | 59.46 | 54.05 |
| **3-motifs ($X^2$)** | 86.49 | **97.30** | 71.62 | 71.62 | 71.62 |
| **4-motifs ($X^2$)** | 89.19 | **94.59** | 64.86 | 62.16 | 67.57 |
| **Bipartiveness (ED)** | 45.95 | **51.35** | 37.84 | 37.84 | 40.54 |
| **Odd-cycles motifs of** | 75.68 | **86.49** | 56.76 | 54.05 | 54.05 |
| **length ($l$) =**<br>**3,5,7 ($X^2$)** |  |  |  |  |  |
| **Ensemble:**<br>**Communities,**<br>**3-motifs , 4-**<br>**motifs, odd-**<br>**cycles of**<br>**length ($l$) =**<br>**3,5,7** | 94.59 | **100** | 81.08 | 72.97 | 75.67 |

## 4. Discussion

This study investigates the properties of graph Laplacian spectrum, the multiscale topological descriptors of the networks and their association over dMRI-based structural brain networks (SBN). All the aforementioned estimators and associations were explored for their repeatability (test-retest scans), their individuality (brain fingerprinting) and how they could be affected by the choice of graph construction scheme. Finally, the evidences of this study were supported statistically but also through perturbations by comparing original values with surrogates ones estimated over random null network models (edge rewiring). For that purpose, I adopted and analyzed the test-retest diffusion-MRI data set from the multimodal neuroimaging database of the Human Connectome Project (HCP) (Glasser et al., 2013; S N Sotiropoulos et al., 2013; Van Essen et al., 2013).

Graph laplacian spectrum of SBN has been studied as a whole but also in sub-ranges of low, middle and large eigenvalues. Every Laplacian eigenvalues subrange has been associated with specific topological patterns covering various spatial scales. The smallest eigenvalues of the Laplacian spectrum reflect the modular organization of a network (Donetti, 2005; Fortunato, 2010; Shen and Cheng, 2010; Shi and Malik, 2000). Repeated duplications and additions of nodes and motifs in the construction of a network leave traces in the network’s Laplacian spectrum in the middle eigenvalues (Banerjee and Jost, 2009, 2008). De Lange et al., (2016) revealed that global symmetry shaped neural spectra and the overlap in the wiring pattern of brain regions measured with MIN can explain the large central peak observed in spectra of neural networks (λ=1). The largest eigenvalue of the Laplacian spectrum informs us of the level of ‘bipartiteness’ of the most bipartite subpart of the network, which is closely related to the number of odd cyclic motifs in the network (Bauer and Jost, 2009). A recent study on SBN introduced a new framework that placed integration and segregation on the end of a continuum of structural connectivity graph Laplacian harmonics via the presentation of a gap-spectrum (Sipes et al., 2024). Their approach partitions graph Laplacian spectrum into integrative, segregative, and degenerate harmonics. It is important to underline here that the whole analysis focused on normalized Laplacian spectrum.

This study showed that normalized Laplacian eigenvalues of dMRI-based structural brain networks are subject-specific, and therefore be used to ‘fingerprint’ an individual with absolute accuracy (100%). Normalized Laplacian eigenvalues are also repeatable across the five graph construction schemes but their connectome-related information of the studying SBN is highly dependent on the graph construction scheme.

The repeatability of Laplacian eigenvectors (connectome harmonics; Naze et al., 2021) is highly dependent on the graph construction scheme. 9m-OMST graph construction scheme showed the smallest group-mean D^LS^ followed by the NS-OMST but without reaching the significant level (p < 0.05, Bonferroni corrected). These findings are supported also by the direct comparison of original MI values with the surrogate D^LS^ values. However, the group-mean D^LS^ even for the 9m-OMST method is far away from characterized as repeatable.

Investigation of the small eigenvalues from the Laplacian spectrum untangled a community structure for the first two graph construction schemes (NS-OMST, 9m-OMST). For those graph construction schemes, the optimal division of the network based on the eigen-difference is suggested to be between 7 and 9 communities (Dimitriadis et al., 2021). My findings are supported by the direct comparison with the surrogate null models. In a recent exploratory study on the same dataset comparing thirty-three graph partition schemes and the same set of graph construction schemes, we revealed a consensus set of 9 communities (Dimitriadis et al., 2021; Fig.5). My analysis showed that the K-Means clustering applied over the first eigenvectors is a better approach compared to the eigen-difference (eigen-gap) method applied over Laplacian eigenvalues. For the first two graph construction schemes (NS-OMST, 9m-OMST), the MI values for the K-Means algorithm demonstrated a high repeatability between the two scans, while the communities extracted with K-Means applied over SBN constructed with 9m-OMST method showed a high similarity with our previous study (*MI* = 0.91 ± 0.04) where a large number of graph partition algorithms were applied on the same set (Dimitriadis et al., 2021).

The Laplacian spectrum of the graph construction schemes showed a clear smooth peak at the middle range as is shown in Fig.3. This peak around λ = 1 suggests a number of motif duplications where its height is relevant to this number. The Laplacian spectrum didn’t show any other clear peaks indicative of recurrent addition of motifs. The observation of this peak on the middle subpart of the Laplacian spectrum was consistent across the graph construction schemes while it was less peaked (lower amplitude) in the rewired surrogate networks. I explored a possible relationship between the RF extracted from the Laplacian spectrum linked to λ = 1 and the total number of 3,4-motifs extracted from the SBN plus the MIN. The adopted multilinear regression analysis untangled a significant trend between the RF and the 3,4-motifs plus the MIN only for the 9m-OMST graph construction scheme which was repeatable. These findings complement to the results of De Lange et al., (2016) showing that local recursive topological patterns expressed here with 3,4-motifs and global symmetry measured with MIN shaped graph Laplacian spectrum explaining partly the RF related to the large central peak observed in the Laplacian spectrum of SBN (λ = 1).

Complementary to the aforementioned analysis, I investigated how nodal weighted clustering coefficient is correlated to the nodal 3-motifs distribution. A positive correlation between the second 3-motif and weighted clustering coefficient was consistently observed across graph construction schemes while a mixed sign of correlation was detected for the first 3-motif with the five graph construction schemes. Correlation between individual distributions of 3,4-motifs across graph construction schemes revealed a strong positive correlation between the 9-m-OMST, NS-thr and NS-t/MD-w. This means that the individual SBN share a large number of common local topologies that is reflected to the global network level.

The largest Laplacian eigenvalues reflect the level of bipartiteness of the structural brain networks. Bipartiteness is related to the odd cyclic motifs, and especially is linked to the triangle motifs and high clustering coefficient observed in small-world brain networks (Bassett and Bullmore, 2006; Bullmore and Sporns, 2009; Hagmann et al., 2008; van den Heuvel et al., 2008). In the present study, bipartiteness which is linked to the largest Laplacian eigenvalue Ln was significantly different on the original SBN compared to the surrogate null networks only for the two graph construction schemes (NS-OMST, 9m-OMST). The exhaustive quantification of odd-cycles with length = 3,5 and 7 over original SBN differed significantly from the ones estimated over surrogate null models for every graph construction scheme. Multi-linear regression analysis between the λn and the bs plus the three total number of odd-cycles (length = 3,5,7) revealed interesting findings only for the two graph construction schemes (NS-OMST, 9m-OMST). Especially for the 9m-OMST, the outcome of this analysis revealed a significant model with a high R^2^ = 0.844 that involved every independent variable, the bs and the three odd-cycles. It is the very study that revealed such a trend between the λn, the bs and the odd-cycles in network science and especially in SBN.

The repeatability of harmonics’ partitions with the gap-spectrum method is higher for the integrative harmonic, middle for the degenerate and lower for the segregative regime. Group-averaged ELD values for every regime of harmonics and across graph construction schemes were significantly different compared to the surrogate ELD values (p < 0.01 x 10^-5^, Bonferroni corrected). Statistical analysis between the three regimes of harmonics in a pairwise fashion and within every graph construction scheme showed significant differences for every pair and graph construction scheme (p < 0.01 x 10^-12^, Bonferroni corrected). No significant difference was detected across the graph construction schemes for integrative and degenerate harmonics while group-averaged ELD was statistically lower for segregative harmonics for 9-m-OMST graph construction scheme compared to the rest of graph construction schemes (p < 0.02 x 10^-12^, Bonferroni corrected).

Under the brain fingerprinting framework, the Laplacian eigenvalues and the topologies of SBN showed an absolute accuracy (100%). Laplacian eigenvectors reached an absolute accuracy only for the 9m-OMST graph construction scheme. In parallel, the performance of brain fingerprinting employing five studying structural properties revealed the communities, and 3,4-motifs across the five construction schemes succeeding the highest performance for 9m-OMST (B), and NS-OMST (A) with the former to get higher accuracies. The combination of the outcome of brain identification strategy for communities, 3,4-motifs and odd-cycles of length (*l*) = 3,5,7 (ensemble way) gave an absolute accuracy (100%) for the 9m-OMST (B), and a 94.59 for the NS-OMST (A). These brain fingerprinting findings support the individuality of the SBN, of the Laplacian spectrum and of the multi-scale resolution of network topology extended from 3,4-motifs and odd-cycles of length (*l*) = 3,5,7 up to communities. The outcome of alternative inputs on brain fingerprinting approach differed across graph construction schemes while 9m-OMST demonstrated the highest performance followed by the NS-OMST.

In this study, I examined the relationship between Laplacian spectrum and topological patterns of SBN quantified by the exhaustive quantification of recursive motifs and odd cycles. The RF linked to λ = 1 was partially predicted by the total number of 3,4-motifs and the MIN only for the 9m-OMST and in both scans. Similarly, the λn was largely predicted by the total number of odd-cycles of length (*l*) = 3,5,7 and the bs for the 9m-OMST and to a less extent for the NS-OMST. These relationships between Laplacian spectrum and network topologies revealed to what extent recursive topological motif and odd-cycles, MIN and bs shaped graph Laplacian spectrum. Although graph Laplacian spectrum properties can now be captured by recursive topological motifs up to some extent, the graph Laplacian spectrum can reveal systems-level changes that cannot be described or detected by standard network metrics.

I investigated also the repeatability and the influence of graph construction schemes on basic Laplacian properties apart from the three sub-ranges of Laplacian spectrum. The range of Synchronizability showed a higher dependency on the graph construction scheme compared to the Laplacian energy. The smallest group-mean between-scan difference for Synchronizability, and for Laplacian energy was shown for the 9m-OMST and NS-OMST graph construction schemes. Both observations were supported statistically by a direct comparison with the corresponding values estimated from the surrogate null networks.

The spatial resolution of structural brain networks restricted by the adopted anatomical atlas, is likely to have a high impact on the network topology and the relevant Laplacian spectrum (Bullmore and Sporns, 2012, 2009). My findings can be considered only for reconstructed anatomical brain networks based on dMRI (Hagmann et al., 2008; Iturria-Medina et al., 2008; van den Heuvel and Hulshoff Pol, 2010), the data-acquisition parameters, the algorithm performing the tractography analysis with its parameters, the atlas template (AAL) and the graph construction scheme. The combination of these choices across the whole analysis might alter the constructed SBN which in turn have impact on the shape of graph Laplacian spectrum and any of the recursive topological motif and odd-cycles (Chung et al., 2003; van den Heuvel et al., 2016).

The Laplacian Spectrum of functional brain networks using electroencephalography, magnetoencephalography, and functional magnetic resonance imaging (fMRI) might differ from structural brain networks from dMRI. Future studies should reveal the relationship between the Laplacian spectrum of functional and structural brain networks. Recent studies used atlas-free connectome harmonics dMRI as a dependent variable to predict the brain resting-state activity of fMRI (independent variables). They showed their findings in datasets across the landscape of consciousness with very interesting findings (Atasoy et al., 2018a, 2016; Luppi et al., 2020) and in psychedelic (Atasoy et al., 2018b) while they explored the robustness of connectome harmonics using local gray matter and long-range white matter (Naze et al., 2021). It would be very important to explore the repeatability of the Laplacian spectrum on structural brain networks from dMRI in an atlas-free scenario.

The structural brain networks microscopically showed a community structure as it has been observed in other species and other types of networks (de Lange et al., 2014). Macroscopically, anatomical connectivity topology is shaped by evolutionary growth constraints that attempt to balance the optimal efficiency and robustness of the communication of various brain networks while simultaneously minimizing wiring cost (Bullmore and Sporns, 2012; Collin et al., 2014; van den Heuvel and Sporns, 2013a, 2013b). The 9m-OMST graph construction scheme that integrates nine diffusion metric-based structural brain networks into one has at its core the OMST topological filtering methodology that optimizes efficiency routing via wiring cost (Dimitriadis et al., 2018, 2017b, 2017c, 2017a; Messaritaki et al., 2019).

### Limitations

My study had a few limitations which is important to discuss. My results were extracted by analysing the HCP dataset, and for that reason it cannot be generalized to other datasets that are acquired with different protocols or scanners or analytic pipelines involving alternative tractography algorithms. It is highly recommended to every researcher to record a percentage of the original cohort across three or more scanning sessions. Moreover, scanning the subjects at the same time of day would be also desirable (Trefler et al., 2016). Many alternative topological filtering schemes have been proposed so far. In my study, I fixed the sparsity of the thresholded networks such as to have the same sparsity as the OMST networks. Other alternative topological filtering schemes with arbitrary or data-driven approach could be also explored under the reproducibility framework. It is important to note here that many other variables can affect the repeatability of structural brain network analyses. There variables include: the parcellation scheme used, the time interval between the test-retest scans, and the resolution of the MR data. There variables should be considered when interpreting structural brain network studies, and a useful discussion of this subject is provided by Welton et al. (2015). It is important to underline here, that I investigated high-order interactions focusing on 3,4-motifs, and odd cycles derived from SBN. There are also alternatively high-order interactions like hypergraphs, and simplicial complexes (Battiston et al., 2020,2021) that weren’t explored here. It is important to underline here, that Interdependencies between brain areas can be explored either from anatomical (structural) perspective (structural connectivity) or by considering statistical interdependencies (functional connectivity). Structural connectivity is typically pairwise, where white-matter fiber tracts start in a certain region, and arrive at another brain area. So, by construction SBN tabulates pairwise associations compared to functional brain networks, which can be built upon high-order interactions (Herzog et al., 2024).

## 5. Conclusions

This study investigates the properties of graph Laplacian spectrum, the multiscale recursive topological patterns of the networks and their association over dMRI-based structural brain networks (SBN). All the aforementioned estimators and associations were explored for their repeatability (test-retest scans), their individuality (brain fingerprinting) and how they could be affected by the choice of graph construction scheme. Finally, the evidences of this study were supported statistically but also through perturbations by comparing original values with surrogates ones estimated over random null network models (edge rewiring). Further analysis is needed by adopting a Multishell Diffusion MRI-Based Tractography, an atlas of higher resolution and a test-retest dataset from another site to evaluate the main findings of this study.

## Acknowledgements

We are grateful to the Human Connectome Project for making the test-retest data freely available.

Data were provided [in part] by the Human Connectome Project, WU-Minn Consortium (Principal Investigators: David Van Essen and Kamil Ugurbil; 1U54MH091657) funded by the 16 NIH Institutes and Centers that support the NIH Blueprint for Neuroscience Research; and by the McDonnell Center for Systems Neuroscience at Washington University

SID is supported by a Beatriu de Pinós fellowship (2020 BP 00116). BP programme is co-funded by the European Union through the COFUND programme (contract number 801370) for Marie-Skłodowska-Curie actions in the “Horizon 2020” programme. The BP programme is geographically localized in Catalonia (Spain) and is driven by AGAUR.

## AUTHOR CONTRIBUTIONS

**Stavros I. Dimitriadis:** conceptualization, methodology, software, validation, formal analysis, investigation, data curation, roles/writing - original draft and funding acquisition.

## Data and code availability

The HCP test-retest data is freely available as listed above.

- **Probabilistic tractography** has been realized with MRtrix (https://www.mrtrix.org/)
- **Network construction** using ExploreDTI-4.8.6 (http://www.exploredti.com/) The code used to generate the graphs for the structural brain networks with the OMST schemes is available at: https://github.com/stdimitr/multi-group-analysis-OMST-GDD.
- **Brain Connectivity Toolbox** (https://sites.google.com/site/bctnet/) The construction of Surrogate null models and the estimation of weighted clustering coefficient and 3,4-motifs has been done with the following equations **Surrogate null models**: randmio_und_connected with iter = 10, **Weighted clustering coefficient** : clustering_coef_wu.m, **3,4-motifs**: motif3fstruct_wei.m; motif4struct_wei. **Matching Index (MIN)**: matching_index_und.m
- **3-odd-cycles : MATLAB :** cycles = allcycles(G,**’MaxCycleLength’**,l); % l = 3,5,7
- **Integrative, segregative, and degenerate harmonics of the structural connectome** The estimation of gap-spectrum for the definition of the three regimes of harmonics was realized with the following equation: **Gap Spectrum Estimation** : find_ev_IntDegSeg.m https://github.com/Raj-Lab-UCSF/IntDegSeg/tree/main
- **Multiscale Graph Comparison via the Embedded Laplacian Discrepancy (ELD)** ELD metric is presented by the authors on the following github page implemented in python (ELD.py) https://github.com/edrictam/Embedded-Laplacian-Distance
- I implemented **ELD in MATLAB** in my personal github webpage based on author’s definition in conjunction to the estimation of graph Laplacian spectrum and Brain Fingerprinting approach.
- **Violin Plots (Figures 9 and 14):** https://zenodo.org/records/12749045 Povilas Karvelis (2025). daviolinplot - violin and raincloud plots (https://github.com/povilaskarvelis/DataViz/releases/tag/v3.2.7), GitHub. Retrieved March 16, 2025.
- **Correlograms (Figures 10 and 11):** SerhanYilmaz(2025). Correlogram (https://www.mathworks.com/matlabcentral/fileexchange/133812-correlogram), MATLAB Central File Exchange. Retrieved March 14, 2025.
- **Portrait Divergence (PID):** An information-theoretic, all-scales approach to comparing networks https://github.com/bagrow/network-portrait-divergence
- **Embedded_Laplacian_Discrepancy (ELD)** ELD is implemented in Python in the author’s website. https://github.com/edrictam/Embedded-Laplacian-Distance A MATLAB implementation of the ELD is provided in my personal github’s website in a repository dedicated to this study (see below).
- A list of MATLAB functions used in the current study is demonstrated in: https://github.com/stdimitr/graph_laplacian_dMRI_repeat_scans

## Abbreviations

MI: mutual information
ED: Euclidean Distance
dMRI: diffusion magnetic resonance imaging
Pcc: Pearson’s correlation coefficient
OMST: orthogonal-minimal-spanning-tree
SBN: structural brain network
AAL: Automated Anatomical Labeling
iFOD2: Second-order Integration over Fiber Orientation Distributions
GFA: general fractional anisotropy
ICVF: Intra-Cellular Volume Fraction
ODI: Orientation Dispersion Index
CSD: constrained spherical deconvolution
NS: the number of streamlines
FA: fractional anisotropy
MD: mean diffusivity
RF: relative frequency
SLD: streamline density
DTI: diffusion tensor imaging
DWI: diffusion-weighting images
ELD: Embedded Laplacian Discrepancy
MIN: Matching Index

## Bibliography

Abdelnour, F., Voss, H. U. & Raj, A. Network diffusion accurately models the relationship between structural and functional brain connectivity networks. NeuroImage 90, 335–347 (2014).

Amico E, J. Goñi.The quest for identifiability in human functional connectomes Sci. Rep., 8 (2018), p. 8254

Anderson P.W. More is different Science, 177 (4047) (1972), pp. 393–396

Atasoy, S., Deco, G., Kringelbach, M.L., Pearson, J., 2018a. Harmonic brain modes: A unifying framework for linking space and time in brain dynamics. Neuroscientist 24, 277–293. doi:10.1177/1073858417728032

Atasoy, S., Donnelly, I., Pearson, J., 2016. Human brain networks function in connectome-specific harmonic waves. Nat. Commun. 7, 10340. doi:10.1038/ncomms10340

Atasoy, S., Roseman, L., Kaelen, M., Kringelbach, M.L., Deco, G., Carhart-Harris, R.L., 2017. Connectome-harmonic decomposition of human brain activity reveals dynamical repertoire re-organization under LSD. Sci. Rep. 7, 17661. doi:10.1038/s41598-017-17546-0

Atasoy, S., Vohryzek, J., Deco, G., Carhart-Harris, R.L., Kringelbach, M.L., 2018b. Common neural signatures of psychedelics: Frequency-specific energy changes and repertoire expansion revealed using connectome-harmonic decomposition. Prog. Brain Res. 242, 97–120. doi:10.1016/bs.pbr.2018.08.009

Atay, F.M., Bıyıkoğlu, T., Jost, J., 2006. Network synchronization: Spectral versus statistical properties. Physica D: Nonlinear Phenomena 224, 35–41. doi:10.1016/j.physd.2006.09.018

Bagrow, J. P. & Bollt, E. M. An information-theoretic, all-scales approach to comparing networks. Appl. Netw. Sci. 4, 45 (2019).

Banerjee, A., Jost, J., 2007. Spectral plots and the representation and interpretation of biological data. Theory Biosci. 126, 15–21. doi:10.1007/s12064-007-0005-9

Banerjee, A., Jost, J., 2008. On the spectrum of the normalized graph Laplacian. Linear Algebra Appl. 428, 3015–3022. doi:10.1016/j.laa.2008.01.029

Banerjee, A., Jost, J., 2009. Graph spectra as a systematic tool in computational biology. Discrete Applied Mathematics 157, 2425–2431. doi:10.1016/j.dam.2008.06.033

Banerjee, A., 2012. Structural distance and evolutionary relationship of networks. BioSystems 107, 186–196. doi:10.1016/j.biosystems.2011.11.004

Barabási A.-L., Albert R. Emergence of scaling in random networks. Science, 286 (5439) (1999), pp. 509–512

Barabási A.-L.The network takeover. Nat. Phys., 8 (1) (2011), p. 14–16

Bassett, D.S., Bullmore, E., 2006. Small-world brain networks. Neuroscientist 12, 512–523. doi:10.1177/1073858406293182

Bassett DS, Brown JA, Deshpande V, Carlson JM, Grafton ST, Conserved.variable architecture of human white matter connectivity. NeuroImage2011;54:1262–79.

Bastiani M, Shah NJ, Goebel R, Roebroeck A. Human cortical connectome reconstruction from diffusion weighted MRI: the effect of tractography algorithm. NeuroImage 2012;62:1732–49.

Battiston F, Cencetti G, Iacopini I, Latora V, Lucas M, Patania A, Young JG, Petri G. Networks beyond pairwise interactions: Structure and dynamics. Physics Reports,Volume 874,2020,Pages 1-92,10.1016/j.physrep.2020.05.004.

Battiston, F., Amico, E., Barrat, A. et al. The physics of higher-order interactions in complex systems. Nat. Phys. 17, 1093–1098 (2021). 10.1038/s41567-021-01371-4

Bauer F, Jost J. Bipartite and neighborhood graphs and the spectrum of the normalized graph Laplacian. Comm. Anal. Geom. 21 (2013), no. 4, 787–845

Belkin M and Niyogi P. 2003. Laplacian eigenmaps for dimensionality reduction and data representation. Neural computation 15, 6 (2003),1373–1396.

Biswas AB, Talya Eden, and Ronitt Rubinfeld. Towards a decomposition-optimal algorithm for counting and sampling arbitrary motifs in sub-linear time. In Approximation, Randomization, and Combinatorial Optimization. Algorithms and Techniques (APPROX/RANDOM), 2021

Boccaletti, S., Latora, V., Chavez, M., Hwang, D.U., 2006. Complex networks: Structure and dynamics. Physics Reports 424, 175–308. doi:10.1016/j.physrep.2005.10.009

Bonacich, P., 1972. Factoring and weighting approaches to status scores and clique identification. J. Math. Sociol. 2, 113–120. doi:10.1080/0022250X.1972.9989806

Bonacich, P., 2007. Some unique properties of eigenvector centrality. Soc. Networks 29, 555–564. doi:10.1016/j.socnet.2007.04.002

Buchanan CR, Pernet CR, Gorgolewski KJ, Storkey AJ, Bastin ME. Test–retest reliabilityof structural brain networks from diffusion MRI. NeuroImage 2014;86:231–43.

Bullmore, E., Sporns, O., 2009. Erratum: Complex brain networks: graph theoretical analysis of structural and functional systems. Nat. Rev. Neurosci. 10, 312–312. doi:10.1038/nrn2618

Bullmore, E., Sporns, O., 2012. The economy of brain network organization. Nat. Rev. Neurosci. 13, 336–349. doi:10.1038/nrn3214

Cheng, X.-Q., Shen, H.-W., 2010. Uncovering the community structure associated with the diffusion dynamics on networks. J. Stat. Mech. 2010, P04024. doi:10.1088/1742-5468/2010/04/P04024

Chung, F.R.K., 1996. Spectral Graph Theory (CBMS Regional Conference Series in Mathematics, No. 92). American Mathematical Society. 1–228.

Collin, G., Sporns, O., Mandl, R.C.W., van den Heuvel, M.P., 2014. Structural and functional aspects relating to cost and benefit of rich club organization in the human cerebral cortex. Cereb. Cortex 24, 2258–2267. doi:10.1093/cercor/bht064

Deco, G., Tononi, G., Boly, M. & Kringelbach, M. L. Rethinking segregation and integration: contributions of whole-brain modelling. Nat. Rev. Neurosci. 16, 430–439 (2015).

Demuru M, M. Fraschini.EEG fingerprinting: subject-specific signature based on the aperiodic component of power spectrum. Comput. Biol. Med., 120 (2020), Article 103748

Demuru M, A.A. Gouw, A. Hillebrand, C.J. Stam, B.W. van Dijk, P. Scheltens,…, P.J. Visser. Functional and effective whole brain connectivity using magnetoencephalography to identify monozygotic twin pairs Sci. Rep., 7 (2017), p. 9685

Desikan, R.S., Ségonne, F., Fischl, B., Quinn, B.T., Dickerson, B.C., Blacker, D., Buckner, R.L., Dale, A.M., Maguire, R.P., Hyman, B.T., Albert, M.S., Killiany, R.J., 2006. An automated labeling system for subdividing the human cerebral cortex on MRI scans into gyral based regions of interest. Neuroimage 31, 968– 980. doi:10.1016/j.neuroimage.2006.01.021

Deslauriers-Gauthier, S., Zucchelli, M., Frigo, M. & Deriche, R. A unified framework for multimodal structure-function mapping based on eigenmodes. Med. Image Anal. 66, 101799 (2020).

Destrieux, C., Fischl, B., Dale, A., Halgren, E., 2010. Automatic parcellation of human cortical gyri and sulci using standard anatomical nomenclature. Neuroimage 53, 1–15. doi:10.1016/j.neuroimage.2010.06.010

Dimitriadis, S.I., Antonakakis, M., Simos, P., Fletcher, J.M., Papanicolaou, A.C., 2017a. Data-Driven Topological Filtering Based on Orthogonal Minimal Spanning Trees: Application to Multigroup Magnetoencephalography Resting-State Connectivity. Brain Connect. 7, 661–670. doi:10.1089/brain.2017.0512

Dimitriadis, S.I., Drakesmith, M., Bells, S., Parker, G.D., Linden, D.E., Jones, D.K., 2017b. Improving the Reliability of Network Metrics in Structural Brain Networks by Integrating Different Network Weighting Strategies into a Single Graph. Front. Neurosci. 11, 694. doi:10.3389/fnins.2017.00694

Dimitriadis, S.I., Messaritaki, E., Jones, D.K., 2021. The impact of graph construction scheme and community detection algorithm on the repeatability of community and hub identification in structural brain networks. Human Brain Mapping 42(13), 4261–4280

Dimitriadis, S.I., Routley, B., Linden, D.E., Singh, K.D., 2018. Reliability of Static and Dynamic Network Metrics in the Resting-State: A MEG-Beamformed Connectivity Analysis. Front. Neurosci. 12, 506. doi:10.3389/fnins.2018.00506

Dimitriadis, S.I., Salis, C., Tarnanas, I., Linden, D.E., 2017c. Topological Filtering of Dynamic Functional Brain Networks Unfolds Informative Chronnectomics: A Novel Data-Driven Thresholding Scheme Based on Orthogonal Minimal Spanning Trees (OMSTs). Front. Neuroinformatics 11, 28. doi:10.3389/fninf.2017.00028

Donetti, L., 2005. Improved spectral algorithm for the detection of network communities, in: AIP Conference Proceedings. Presented at the MODELING COOPERATIVE BEHAVIOR IN THE SOCIAL SCIENCES, AIP, pp. 104–107. doi:10.1063/1.2008598

Estrada, E. The many facets of the Estrada indices of graphs and networks. SeMA 79, 57–125 (2022). 10.1007/s40324-021-00275-w

Farwell LA. Brain fingerprinting: a comprehensive tutorial review of detection of concealed information with event-related brain potentials. Cogn Neurodyn. 2012 Apr;6(2):115–54. doi: 10.1007/s11571-012-9192-2.

Feinberg, D.A., Moeller, S., Smith, S.M., Auerbach, E., Ramanna, S., Gunther, M., Glasser, M.F., Miller, K.L., Ugurbil, K., Yacoub, E., 2010. Multiplexed echo planar imaging for sub-second whole brain FMRI and fast diffusion imaging. PLoS ONE 5, e15710. doi:10.1371/journal.pone.0015710

Fernandes BS, L.M. Williams, J. Steiner, M. Leboyer, A.F. Carvalho, M. Berk The new field of ‘precision psychiatry. BMC Med., 15 (2017), p. 80

Finn ES, X. Shen, D. Scheinost, M.D. Rosenberg, J. Huang, M.M. Chun,…, R.T. Constable Functional connectome fingerprinting: identifying individuals based on patterns of brain connectivity. Nat. Neurosci., 18 (2015), pp. 1664–1671

Fornito A, A. Arnatkevičiūtė, B.D. Fulcher. Bridging the Gap between Connectome and Transcriptome. Trends Cogn. Sci. (Regul. Ed.), 23 (2019), pp. 34–50

Fortunato, S., 2010. Community detection in graphs. Phys. Rep. 486, 75–174. doi:10.1016/j.physrep.2009.11.002

Friston KJ. Functional and effective connectivity: a review. Brain Connect. 2011;1(1):13–36. doi: 10.1089/brain.2011.0008.

Glasser, M.F., Sotiropoulos, S.N., Wilson, J.A., Coalson, T.S., Fischl, B., Andersson, J.L., Xu, J., Jbabdi, S., Webster, M., Polimeni, J.R., Van Essen, D.C., Jenkinson, M., WU-Minn HCP Consortium, 2013. The minimal preprocessing pipelines for the Human Connectome Project. Neuroimage 80, 105–124. doi:10.1016/j.neuroimage.2013.04.127

Goemans MX. Lecture notes on bipartite matching. MIT. 18.453: Combinatorial Optimization https://math.mit.edu/~goemans/18433S09/matching-notes.pdf

Griffa, A., Amico, E., Liégeois, R., Van De Ville, D., Preti, M.G., 2022. Brain structure-function coupling provides signatures for task decoding and individual fingerprinting. Neuroimage 250, 118970. doi:10.1016/j.neuroimage.2022.118970

Hagmann, P., Cammoun, L., Gigandet, X., Meuli, R., Honey, C.J., Wedeen, V.J., Sporns, O., 2008. Mapping the structural core of human cerebral cortex. PLoS Biol. 6, e159. doi:10.1371/journal.pbio.0060159

Hagmann P, Cammoun L, Gigandet X, Meuli R, Honey CJ, Wedeen VJ, et al. Mapping the structural core of human cerebral cortex. PLoS Biol 2008;6(7):e159.

Hakimi-Nezhaad, M., Ashrafi, A.R., 2014. A note on normalized Laplacian energy of graphs. J. Contemp. Mathemat. Anal. 49, 207–211. doi:10.3103/S106836231405001X

Hampel H, A. Vergallo, G. Perry, S. Lista, Alzheimer Precision Medicine Initiative (APMI) The Alzheimer Precision Medicine Initiative. J. Alzheimer’s Disease: JAD, 68 (2019), pp. 1–24

Harriger, L., van den Heuvel, M.P., Sporns, O., 2012. Rich club organization of macaque cerebral cortex and its role in network communication. PLoS ONE 7, e46497. doi:10.1371/journal.pone.0046497

Herzog, R., Barbey, F.M., Islam, M.N. et al. High-order brain interactions in ketamine during rest and task: a double-blinded cross-over design using portable EEG on male participants. Transl Psychiatry 14, 310 (2024). 10.1038/s41398-024-03029-0

Hilgetag C.C., Kötter R., Stephan K.E., Sporns O. Computational Methods for the Analysis of Brain Connectivity (2002), pp. 295–335, 10.1007/978-1-59259-275-3_14

Huang, W., Bolton, T.A.W., Medaglia, J.D., Bassett, D.S., Ribeiro, A., Van De Ville, D., 2018. A graph signal processing perspective on functional brain imaging. Proc. IEEE 106, 868–885. doi:10.1109/JPROC.2018.2798928

Iturria-Medina, Y., Sotero, R.C., Canales-Rodríguez, E.J., Alemán-Gómez, Y., Melie-García, L., 2008. Studying the human brain anatomical network via diffusion-weighted MRI and Graph Theory. Neuroimage 40, 1064–1076. doi:10.1016/j.neuroimage.2007.10.060

Jones DK, Knösche TR, Turner R. White matter integrity, fiber count, and other fallacies: the do’s and don’ts of diffusion MRI. NeuroImage 2013;73:239–54.

de Lange, S.C., de Reus, M.A., van den Heuvel, M.P., 2014. The Laplacian spectrum of neural networks. Front. Comput. Neurosci. 7, 189. doi:10.3389/fncom.2013.00189

de Lange S.C., Martijn P. van den Heuvel, Marcel A. de Reus, 2016.The role of symmetry in neural networks and their Laplacian spectra, NeuroImage,Volume 141,Pages 357–365,ISSN 1053-8119, 10.1016/j.neuroimage.2016.07.051.

Leemans, A., Jeurissen, B., Sijbers, J., Jones, D.K., 2009. ExploreDTI: a graphical toolbox for processing, analyzing, and visualizing diffusion MR data. ISMRM.

Lehnertz K, Bröhl T, von Wrede R.Epileptic-network-based prediction and control seizures in humans.Neurobiology of Disease,Volume 181,2023,10.1016/j.nbd.2023.106098.

Lemkaddem A, Daducci A, Kunz N, Lazeyras F, Seeck M, Thiran J, et al. Connectivity and tissue microstructural alterations in right and left temporal lobeepilepsy revealed by diffusion spectrum imaging. NeuroImage: Clin 2014;5:349–58

Liang, Z., King, J., Zhang, N., 2011. Uncovering intrinsic connectional architecture of functional networks in awake rat brain. J. Neurosci. 31, 3776–3783. doi:10.1523/JNEUROSCI.4557-10.2011

Lioi, G., Gripon, V., Brahim, A., Rousseau, F. & Farrugia, N. Gradients of connectivity as graph fourier bases of brain activity. Netw. Neurosci. 5, 322–336 (2021).

Lopez, E. T., Minino, R., Liparoti, M., Polverino, A., Romano, A., De Micco, R., Lucidi, F., Tessitore, A., Amico, E., Sorrentino, G., Jirsa, V., & Sorrentino, P. (2023). Fading of brain network fingerprint in Parkinson’s disease predicts motor clinical impairment. Human Brain Mapping, 44, 1239–1250.

Luppi, A.I., Vohryzek, J., Kringelbach, M.L., Mediano, P.A.M., Craig, M.M., Adapa, R., Carhart-Harris, R.L., Roseman, L., Pappas, I., Finoia, P., Williams, G.B., Allanson, J., Pickard, J.D., Menon, D.K., Atasoy, S., Stamatakis, E.A., 2020. Connectome Harmonic Decomposition of Human BrainDynamics Reveals a Landscape of Consciousness. biorxiv.

Luppi, A.I., Gellersen, H.M., Liu, ZQ. et al. Systematic evaluation of fMRI data-processing pipelines for consistent functional connectomics. Nat Commun 15, 4745 (2024). 10.1038/s41467-024-48781-5

Margulies, D. S. et al. Situating the default-mode network along a principal gradient of macroscale cortical organization. Proc. Natl Acad. Sci. 113, 12574–12579 (2016).

Maslov, S., Sneppen, K., 2002. Specificity and stability in topology of protein networks. Science 296, 910–913. doi:10.1126/science.1065103

McGraw, P.N., Menzinger, M., 2008. Laplacian spectra as a diagnostic tool for network structure and dynamics. Phys. Rev. E Stat. Nonlin. Soft Matter Phys. 77, 031102. doi:10.1103/PhysRevE.77.031102

Medaglia, J.D., Huang, W., Karuza, E.A., Kelkar, A., Thompson-Schill, S.L., Ribeiro, A., Bassett, D.S., 2018. Functional Alignment with Anatomical Networks is Associated with Cognitive Flexibility. Nat. Hum. Behav. 2, 156–164. doi:10.1038/s41562-017-0260-9

Messaritaki, E., Dimitriadis, S.I., Jones, D.K., 2019. Optimization of graph construction can significantly increase the power of structural brain network studies. Neuroimage 199, 495–511. doi:10.1016/j.neuroimage.2019.05.052

Milo R., Shen-Orr S., Itzkovitz S., Kashtan N., Chklovskii D., Alon U.Network motifs: Simple building blocks of complex networks. Science, 298 (5594) (2002), pp. 824–827

Miranda-Dominguez O, B.D. Mills, S.D. Carpenter, K.A. Grant, C.D. Kroenke, J.T. Nigg, D.A. Fair Connectotyping: model Based Fingerprinting of the Functional Connectome. PLoS ONE, 9 (2014), Article e111048

Moeller, S., Yacoub, E., Olman, C.A., Auerbach, E., Strupp, J., Harel, N., Uğurbil, K., 2010. Multiband multislice GE-EPI at 7 tesla, with 16-fold acceleration using partial parallel imaging with application to high spatial and temporal whole-brain fMRI. Magn. Reson. Med. 63, 1144–1153. doi:10.1002/mrm.22361

Müller, E. J., Munn, B. R., Aquino, K. M., Shine, J. M. & Robinson, P. A. The music of the hemispheres: Cortical eigenmodes as a physical basis for large-scale brain activity and connectivity patterns. Front. Hum. Neurosci. 16, 1062487 (2022).

Naze, S., Proix, T., Atasoy, S., Kozloski, J.R., 2021. Robustness of connectome harmonics to local gray matter and long-range white matter connectivity changes. Neuroimage 224, 117364. doi:10.1016/j.neuroimage.2020.117364

Newman, M.E.J., 2006. Modularity and community structure in networks. Proc Natl Acad Sci USA 103, 8577–8582. doi:10.1073/pnas.0601602103

Newman, M., 2003. The Structure and Function of Complex Networks. SIAM Rev. 45, 167–256. doi:10.1137/S003614450342480

Nigro N, R. Ricelli R, L. Passamonti, G. Arabia, M. Morelli, R. Nistico and M. Salsone, G. Barbagallo, A. Quattrone. Characterizing structural neural networks in de novo Parkinson disease patients using diffusion tensor imaging. Hum. Brain Mapp., 37 (2016), pp. 4500–4510

Page, L., Brin, S., Rajeev, M., Terry, W., 2001. The PageRank Citation Ranking: Bringing Order to the Web. Technical Report 1, 1999–1966.

Preti, M.G., Van De Ville, D., 2019. Decoupling of brain function from structure reveals regional behavioral specialization in humans. Nat. Commun. 10, 4747. doi:10.1038/s41467-019-12765-7

Qi, S., Meesters, S., Nicolay, K., ter Haar Romeny, B. M., & Ossenblok, P. (2015). The influence of construction methodology on structural brain network measures: a review. Journal of Neuroscience Methods, 253, 170–182. 10.1016/j.jneumeth.2015.06.016

Raj, A., Kuceyeski, A., Weiner, M., 2012. A network diffusion model of disease progression in dementia. Neuron 73, 1204–1215. doi:10.1016/j.neuron.2011.12.040

de Reus MA, M.P. van den Heuvel.The parcellation-based connectome: limitations and extensions. Neuroimage, 80 (2013), pp. 397–404

Rodrigues J, S. de, F.L. Ribeiro, J.R. Sato, R.C. Mesquita, C.E.B. Júnior Identifying individuals using fNIRS-based cortical connectomes. Biomed. Opt. Express, 10 (2019), pp. 2889–2897

Rubinov M, Sporns O.Complex network measures of brain connectivity: Uses and interpretations.NeuroImage,Volume 52, Issue 3,2010,Pages 1059–1069.10.1016/j.neuroimage.2009.10.003.

Rubner, Y., 2000. :{unav). Springer Science and Business Media LLC. doi:10.1023/a:1026543900054

Sareen E, Zahar S, Dimitri Van De Ville, Anubha Gupta, Alessandra Griffa, Enrico Amico. Exploring MEG brain fingerprints: Evaluation, pitfalls, and interpretations.NeuroImage,Volume 240,2021,118331, 10.1016/j.neuroimage.2021.118331.

Sareen E, L. Singh, A. Gupta, R. Verma, G.K. Achary, B. Varkey.Functional Brain Connectivity Analysis in Intellectual Developmental Disorder During Music Perception. IEEE Trans. Neural Sys. Rehabilit. Eng., 28 (2020), pp. 2420–2430

Setsompop, K., Gagoski, B.A., Polimeni, J.R., Witzel, T., Wedeen, V.J., Wald, L.L., 2012. Blipped-controlled aliasing in parallel imaging for simultaneous multislice echo planar imaging with reduced g-factor penalty. Magn. Reson. Med. 67, 1210–1224. doi:10.1002/mrm.23097

Shen, H.-W., Cheng, X.-Q., 2010. Spectral methods for the detection of network community structure: a comparative analysis. J. Stat. Mech. 2010, P10020. doi:10.1088/1742-5468/2010/10/P10020

Shi, J., Malik, J., 2000. Normalized cuts and image segmentation. IEEE Trans. Pattern Anal. Mach. Intell. 22, 888–905. doi:10.1109/34.868688

Sihag, S., Naze, S., Taghdiri, F., Tator, C., Wennberg, R., Mikulis, D., Green, R., Colella, B., Tartaglia, M.C., Kozloski, J.R., 2020. Multimodal Dynamic Brain Connectivity Analysis Based on Graph Signal Processing for Former Athletes with History of Multiple Concussions. IEEE Trans. on Signal and Inf. Process. over Networks 1–1. doi:10.1109/TSIPN.2020.2982765

Sipes, B.S., Nagarajan, S.S. & Raj, A. Integrative, segregative, and degenerate harmonics of the structural connectome. Commun Biol 7, 986 (2024). 10.1038/s42003-024-06669-6

Smith, R. E., Tournier, J.-D., Calamante, F., & Connelly, A. (2012). Anatomically-constrained tractography: Improved diffusion MRI streamlines tractography through effective use of anatomical information. NeuroImage, 62(3), 1924– 1938. 10.1016/j.neuroimage.2012.06.005,

Smith, R. E., Tournier, J.-D., Calamante, F., & Connelly, A. (2015). SIFT2: Enabling dense quantitative assessment of brain white matter connectivity using streamlines tractography. NeuroImage, 119, 338–351. 10.1016/j.neuroimage.2015.06.092

Sorrentino, P., Rucco, R., Lardone, A., Liparoti, M., Troisi Lopez, E., Cavaliere, C., Soricelli, A., Jirsa, V., Sorrentino, G., & Amico, E. (2021). Clinical connectome fingerprints of cognitive decline. NeuroImage, 238, 118253.

Sotiropoulos, Stamatios N, Jbabdi, S., Xu, J., Andersson, J.L., Moeller, S., Auerbach, E.J., Glasser, M.F., Hernandez, M., Sapiro, G., Jenkinson, M., Feinberg, D.A., Yacoub, E., Lenglet, C., Van Essen, D.C., Ugurbil, K., Behrens, T.E.J., WU-Minn HCP Consortium, 2013. Advances in diffusion MRI acquisition and processing in the Human Connectome Project. Neuroimage 80, 125–143. doi:10.1016/j.neuroimage.2013.05.057

Sotiropoulos, S N, Moeller, S., Jbabdi, S., Xu, J., Andersson, J.L., Auerbach, E.J., Yacoub, E., Feinberg, D., Setsompop, K., Wald, L.L., Behrens, T.E.J., Ugurbil, K., Lenglet, C., 2013. Effects of image reconstruction on fiber orientation mapping from multichannel diffusion MRI: reducing the noise floor using SENSE. Magn. Reson. Med. 70, 1682–1689. doi:10.1002/mrm.24623

Sporns, O., and Kötter, R. (2004). Motifs in brain networks. PLoS Biol. 2:e369. doi: 10.1371/journal.pbio.0020369

Sporns O, G. Tononi, R. Kötter. The Human Connectome: a Structural Description of the Human Brain PLoS Comput. Biol., 1 (2005), p. e42

Stam CJ. Modern network science of neurological disorders. Nat Rev Neurosci. 2014 Oct;15(10):683–95. doi: 10.1038/nrn3801. Epub 2014 Sep 4. PMID: 25186238.

Tam E, Dunson D. Multiscale Graph Comparison via the Embedded Laplacian Discrepancy. 10.48550/arXiv.2201.12064

Taylor PN, C.E. Han, J.-C. Schoene-Bake, B. Weber, M. Kaiser. Structural connectivity changes in temporal lobe epilepsy: spatial features contribute more than topological measures. Neuroimage: Clinic, 8 (2015), pp. 322–328

Tournier, J.-D., Calamante, F., Gadian, D.G., Connelly, A., 2004. Direct estimation of the fiber orientation density function from diffusion-weighted MRI data using spherical deconvolution. Neuroimage 23, 1176–1185. doi:10.1016/j.neuroimage.2004.07.037

Tournier, J.-D., Calamante, F., & Connelly, A. (2010). Improved probabilistic streamlines tractography by 2nd order integration over fibre orientation distributions. In Proceedings of the International Society for Magnetic Resonance in Medicine (Vol. 1670). Stockholm, Sweden.

Tournier, J.-D., Smith, R. E., Raffelt, D., Tabbara, R., Dhollander, T., Pietsch, M., Christiaens, D., Jeurissen, B., Yeh, C.-H., & Connelly, A. (2019). MRtrix3: A fast, flexible and open software framework for medical image processing and visualisation. NeuroImage, 202, 116137. 10.1016/j.neuroimage.2019.116137

Trefler A, Neda Sadeghi, Adam G. Thomas, Carlo Pierpaoli, Chris I. Baker, Cibu Thomas, Impact of time-of-day on brain morphometric measures derived from T1-weighted magnetic resonance imaging, NeuroImage, 133, 2016, 41–52,

Tzourio-Mazoyer, N., Landeau, B., Papathanassiou, D., Crivello, F., Etard, O., Delcroix, N., Mazoyer, B., Joliot, M., 2002. Automated anatomical labeling of activations in SPM using a macroscopic anatomical parcellation of the MNI MRI single-subject brain. Neuroimage 15, 273–289. doi:10.1006/nimg.2001.0978

van den Heuvel, M.P., Stam, C.J., Boersma, M., Hulshoff Pol, H.E., 2008. Small-world and scale-free organization of voxel-based resting-state functional connectivity in the human brain. Neuroimage 43, 528–539. doi:10.1016/j.neuroimage.2008.08.010

van den Heuvel, M.P., Hulshoff Pol, H.E., 2010. Specific somatotopic organization of functional connections of the primary motor network during resting state. Hum. Brain Mapp. 31, 631–644. doi:10.1002/hbm.20893

van den Heuvel, M.P., Sporns, O., 2013a. Network hubs in the human brain. Trends Cogn Sci (Regul Ed) 17, 683–696. doi:10.1016/j.tics.2013.09.012

van den Heuvel, M.P., Sporns, O., 2013b. An anatomical substrate for integration among functional networks in human cortex. J. Neurosci. 33, 14489–14500. doi:10.1523/JNEUROSCI.2128-13.2013

van den Heuvel M.P., Bullmore E.T., Sporns O. Comparative connectomics Trends Cogn. Sci. (2016), 10.1016/j.tics.2016.03.001

Van Essen, D.C., Smith, S.M., Barch, D.M., Behrens, T.E.J., Yacoub, E., Ugurbil, K., WU-Minn HCP Consortium, 2013. The WU-Minn Human Connectome Project: an overview. Neuroimage 80, 62–79. doi:10.1016/j.neuroimage.2013.05.041

Varshney, L.R., Chen, B.L., Paniagua, E., Hall, D.H., Chklovskii, D.B., 2011. Structural properties of the Caenorhabditis elegans neuronal network. PLoS Comput. Biol. 7, e1001066. doi:10.1371/journal.pcbi.1001066

Vukadinović, D., Huang, P., Erlebach, T., 2002. On the spectrum and structure of internet topology graphs, in: Unger, H., Böhme, T., Mikler, A. (Eds.), Innovative Internet Computing Systems, Lecture Notes in Computer Science. Springer Berlin Heidelberg, Berlin, Heidelberg, pp. 83–95. doi:10.1007/3-540-48080-3_8

Wang, M.B., Owen, J.P., Mukherjee, P., Raj, A., 2017. Brain network eigenmodes provide a robust and compact representation of the structural connectome in health and disease. PLoS Comput. Biol. 13, e1005550. doi:10.1371/journal.pcbi.1005550

Watts D.J., Strogatz S.H.Collective dynamics of ‘small-world’ networks. Nature, 393 (6684) (1998), p. 440

Welton, T., D.A. Kent, D.P. Auer, R.A. Dineen.Reproducibility of graph-theoretic brain network metrics: a systematic review. Brain Connect., 5 (4) (2015), pp. 193–202

Xu, J., Moeller, S., Strupp, J., Auerbach, E., Chen, L., Feinberg, D.A., Ugurbil, K., Yacoub, E., 2012. Highly accelerated whole brain imaging using aligned-blipped-controlled-aliasing multiband EPI. Proc. Int. Soc. Mag. Reson. Med 20, 2306.

Zalesky A, Fornito A, Harding IH, Cocchi L, Yücel M, Pantelis C, et al. Whole-brain anatomical networks: does the choice of nodes matter? NeuroImage2010;50:970–83.

Zhang Z, Liao W, Chen H, Mantini D, Ding JR, Xu Q, et al. Altered functional– structuralcoupling of large-scale brain networks in idiopathic generalized epilepsy. Brain2011;134:2912–28.

